# A salt-induced kinase (SIK) is required for the metabolic regulation of sleep

**DOI:** 10.1101/586701

**Authors:** Jeremy J. Grubbs, Lindsey E. Lopes, Alexander M. van der Linden, David M. Raizen

## Abstract

Many lines of evidence point to links between sleep regulation and energy homeostasis, but mechanisms underlying these connections are unknown. During *C. elegans* sleep, energetic stores are allocated to non-neural tasks with a resultant drop in the overall fat stores and energy charge. Mutants lacking KIN-29, the *C. elegans* homolog of a mammalian Salt-Inducible Kinase (SIK) that signals sleep pressure, have low ATP levels despite high fat stores, indicating a defective response to cellular energy deficits. Liberating energy stores corrects adiposity and sleep defects of *kin-29* mutants. *kin-29* sleep and energy homeostasis roles map to a small number of sensory neurons that act upstream of fat regulation as well as of central sleep-controlling neurons, suggesting hierarchical somatic/neural interactions regulating sleep and energy homeostasis. Genetic interaction between *kin-29* and the histone deacetylase *hda-4* coupled with subcellular localization studies indicate that KIN-29 acts in the nucleus to regulate sleep. We propose that KIN-29/SIK acts in nuclei of sensory neuroendocrine cells to transduce low cellular energy charge into the mobilization of energy stores, which in turn promotes sleep.

**Highlights:**

- Sleep is associated with fat mobilization and low ATP levels
- Metabolic regulation of sleep requires the salt-induced kinase (SIK) homolog KIN-29
- KIN-29 acts in sensory neurons upstream of sleep-promoting neurons
- Nuclear localization of KIN-29 is required for the metabolic regulation of sleep
- A type 2 histone deacetylase acts down stream of KIN-29 in the regulation of sleep.
- Liberation of energy from fat-storage cells promotes sleep.
- Beta-oxidation promotes sleep.

## INTRODUCTION

Sleep is intricately connected with metabolism, and reciprocal interactions between sleep and metabolic processes underlie a number of clinical pathologies. Acute disruption of human sleep results in elevated appetite [1] and insulin resistance [2], and chronic short sleeping humans are more likely to be obese and diabetic [3]. Starvation in humans, rats, *Drosophila*, and *C. elegans* affects sleep [4-9], indicating that sleep is regulated, in part, by nutrient availability.

Sleep is associated with reduced neural energetic demands across phylogeny [10-12]; for example, slow wave sleep in mammals is associated with reduced nervous system energetic demands [13-15], and the reduction in neural activity in a sleeping *C. elegans* is likely similarly associated with reduced energy demands [11]. Despite this apparent reduced energy demand in neurons, overall metabolic rates during sleep in mammals [15, 16] and *Drosophila* [17] are only modestly reduced, suggesting that during sleep energetic stores are allocated to other metabolic functions [18], such as the synthesis of proteins [19] and other macromolecules [20]. Importantly, while there are genes reported to function both in metabolic regulation and in sleep [21-26], mechanisms by which these gene products couple the two processes at the level of the whole organism remain unclear. Thus, the molecular and cellular mechanisms connecting sleep with energy homeostasis of the animal remain opaque.

Salt-Inducible Kinases (SIKs) have been identified as conserved regulators of sleep [27] and metabolism [28]. There are three SIKs in mammals, two in *Drosophila* and one in *C. elegans* called KIN-29 [29]. Gain-of-function mouse mutants of SIK3 are sleepy [27] with a phosphoprotein profile that mimics that of sleep-deprived mice [30], indicating that SIK3 signaling promotes sleep need. The *Drosophila* SIK3 and *C. elegans* KIN-29 loss-of-function mutants have reduced sleep [27]. *kin-29* is also required for sleep in satiated animals [31, 32], suggesting a generalized role for KIN-29 in promoting sleep.

In addition to sleep behavioral phenotypes, SIK gene mutations are associated with metabolic defects. In *Drosophila*, reduction of dSIK gene function in neurons results in elevated levels of triglycerides and glycogen [33], whereas loss of *Drosophila* SIK3 in the fat body results in a depletion of triglyceride stores [34]. Based on this combination of sleep and metabolic phenotypes, we hypothesized that SIKs function may be an integral part of the mechanism by which sleep and energy homeostasis are integrated. We set out to test this hypothesis using *C. elegans*, which has proven a powerful organism to study sleep [35] as well as metabolism [36].

*C. elegans* sleeps during development in a stage known as developmentally-timed sleep (DTS), or lethargus [37, 38]. They also sleep after exposure to environmental conditions that cause cellular stress in a behavior termed stress-induced sleep (SIS) [39, 40]. Additionally, *C. elegans* sleep when satiated [32, 41] and in the setting of starvation [4, 32]. Two neurons show strong effects in regulating *C. elegans* sleep, the RIS neuron regulates DTS [42], starvation-associated sleep [32], and satiety-associated sleep [32], and the ALA neuron regulates SIS [39, 43, 44].

Here, we show that multiple types of *C. elegans* sleep are associated with reduced energy levels of the animals, and require the function of KIN-29/SIK. Consistent with a deficit in energy mobilization, we show that *kin-29* mutants have reduced ATP and behave like starved animals despite having elevated fat stores. Experimental mobilization of triglycerides in these fat stores restores sleep. While mouse SIK3 acts in the cytoplasm to phosphorylate multiple brain targets [30], we find that *C. elegans kin-29* acts in the nucleus to regulate sleep and energy homeostasis via just a single gene called *hda-4*, which encodes a class IIa histone-deacytelase. KIN-29 and HDA-4 act in the same metabolically-responsive sensory neurons upstream of fat homeostasis and of the activation of the sleep-promoting neurons ALA and RIS. Together, these results indicate that sleep is regulated via hierarchical interactions between neurons that sense energy needs, fat storage cells that respond to energy need, and the action of sleep-inducing neurons.

## RESULTS

### Sleep is associated with energetic store depletion and fat mobilization

To understand how sleep/wake states are coordinated with the metabolic states of an animal, we studied how lethargus (DTS), stress-induced sleep (SIS), and sleep deprivation (SD) correlate with metabolic parameters. We hypothesized that during lethargus/DTS, when *C. elegans* faces the energetically expensive task of replacing its exoskeleton [45], it sleeps to conserve energy [15] and also mobilizes fat to release energy for biosynthetic tasks. Likewise, during conditions of cellular stress, such as after genotoxic ultraviolet light exposure, *C. elegans* sleeps presumably to conserve energy and mobilizes fat to release energy needed for cellular repair. If this hypothesis is correct, then energy levels should be inversely proportional to sleep drive. When energy levels are low or dropping, sleep drive is high, and when energy levels are high or increasing, sleep drive is low.

To assess energy levels, we measured ATP in whole animals. Consistent with our hypothesis and with published data [46], ATP levels built up until L1 lethargus/DTS, dropped during L1 lethargus, and then began to recover after L1 lethargus (**Fig 1A**). Similarly, after exposure to genotoxic stress, ATP levels decreased for four hours, a period in which the animals slept (**Fig 1C**). These data indicate that during SIS and DTS there are high energetic demands and that the rate of ATP consumption exceeds the rate of ATP generation.

**Fig 1.**
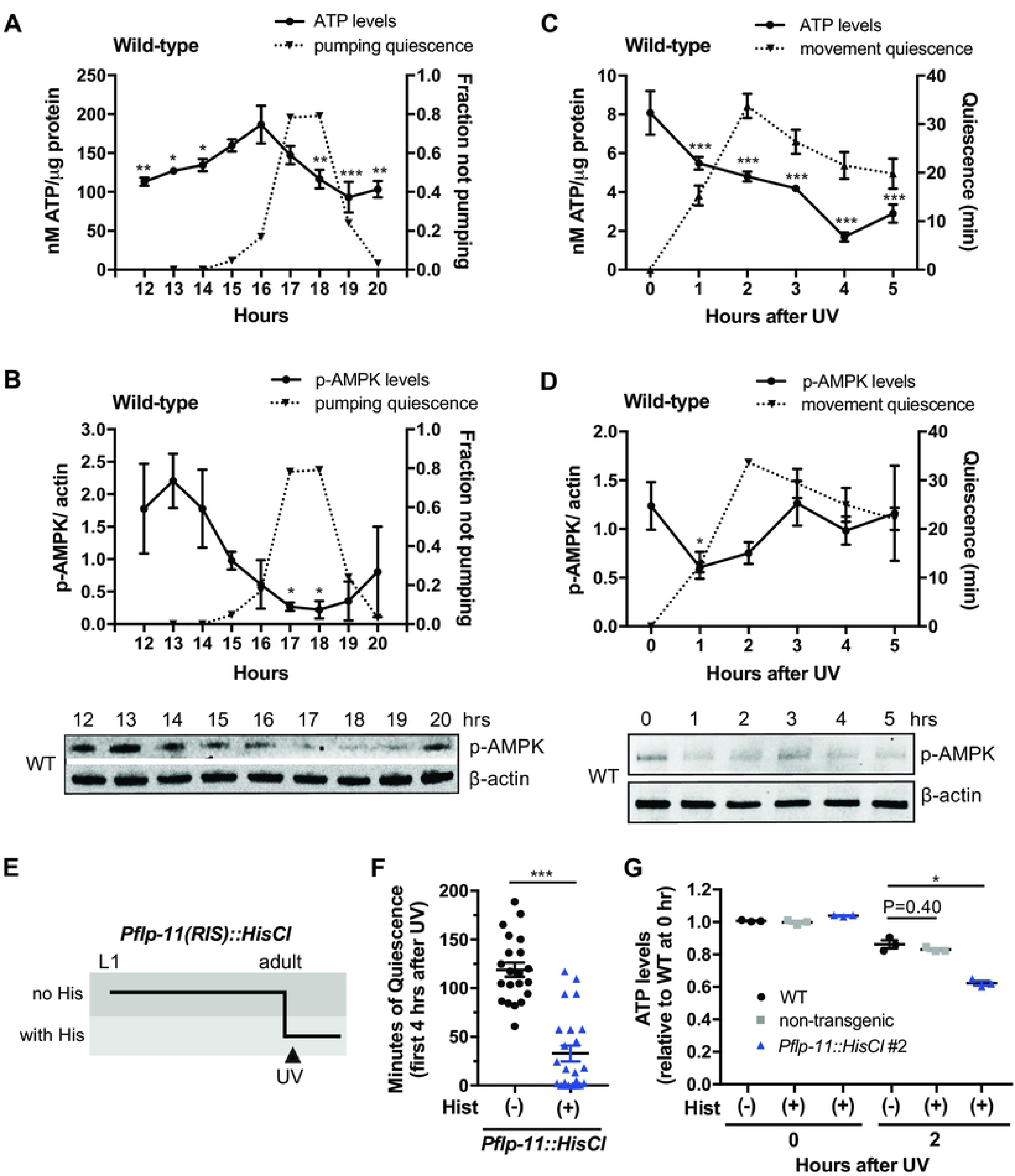
ATP and p-AMPK levels are reduced during DTS and SIS. **(A and B)** Levels of total body ATP (nM ATP per μg protein) (A) and total body normalized phosphorylated AMPK (p-AMPK relative to actin loading control) (B) measured in wild-type animals before, during and after lethargus/DTS of the first larval (L1) stage. Shown is one representative time-course of an experiment run independently seven times for ATP (see **Fig 3D**) and three times for p-AMPK (see **Fig 3E**). p-AMPK and ATP levels were measured from the same samples, and data is represented as the mean ± SD with 2 technical replicates for each. Statistical comparisons were performed with a 2-way ANOVA, followed by post-hoc pairwise comparisons at each time-point to obtain nominal p-values, which were subjected to a Tukey’s multiple comparison correction. ***, ** and * indicates corrected p values that are different from the greatest value in the time-course for DTS at p<0.001, p<0.01 and p<0.05, respectively. A representative Western blot is shown below the graph where the intensity of the upper bands represents levels of p-AMPK, and the intensity of the lower bands represent levels of the loading control β-actin. **(C and D)** Levels of total body ATP (nM ATP per μg protein) (C) and total body normalized p-AMPK (relative to actin loading control) (D) in wild-type adults after UVC irradiation/SIS. Body movement quiescence was measured on the same samples used for ATP measurements. Shown is one representative time-course of an experiment run independently six times for ATP (see **Fig 3H**). Data is represented as the mean ± SD with 3 technical replicates for ATP. For p-AMPK, data are represented as the mean ± SEM of 2 experiments. Statistical comparisons were performed with a 2-way ANOVA, followed by post-hoc pairwise comparisons at each time-point to obtain nominal p-values, which were subjected to a Tukey’s multiple comparison correction. *** and * indicates corrected p values that are different from 0 hours for SIS at p<0.001, and p<0.05, respectively. A representative Western blot of p-AMPK and actin levels is shown below the graph. **(E)** Schematic of a histamine-mediated pharmacogenetic inhibition experiment to deprive worms of stress-induced sleep (SIS). Age-matched worms were grown from the L1 stage to adulthood in the absence of histamine. One-day old adults were transferred to individual wells of a WorMotel device with each well containing 10 mM histamine or vehicle control agar. Approximately 15 min after transfer, worms were exposed to UVC irradiation (1,500 J/m^2^) and their body movement quiescence was recorded for 8 hours. Histamine causes a hyperpolarization. See also **Material and Methods**. **(F)** Minutes of body movement quiescence during the first 4 hours after UVC exposure/SIS in adult animals expressing either the *Pflp-11::HisCl* transgene (+) or in non-transgenic animals (-). The *flp-11* promoter is expressed in RIS. Shown is one representative replicate of an experiment run three times. Data are represented as mean ± SEM. *** p<0.001, Student’s two-tailed *t* test. **(G)** Levels of total body ATP (nM ATP/μg protein) measured 0 and 2 hours (during maximal movement quiescence) after UVC exposure in adult animals expressing either the *Pflp-11::HisCl* transgene or in non-transgenic animals in the presence of 10 mM histamine (+His) compared to wild-type (WT) adults in the absence of histamine (-His). The *flp-11* promoter is expressed in RIS. Data were normalized to wild-type controls immediately before UVC exposure (0 hours) in the absence of histamine (-His). The graph shows the mean ± SEM of 3 experiments. * indicate values that are different from that of non-transgenic animals (+His) at p<0.05, Student’s two-tailed *t* test.

Adenosine monophosphate regulated protein kinase (AMPK) is a conserved regulator of metabolism and energy at the cellular and whole body level [47]. AMPK is activated by a high ratio of AMP to ATP; its activation inhibits anabolic pathways and activates catabolic pathways [47]. We measured activation of the *C. elegans* AMPKα2 homolog AAK-2 using an antibody directed against the phosphorylated threonine 171 of mammals AMPK, which is equivalent to threonine 243 in *C. elegans* AAK-2 (**S1B Fig**). This anti-p-AMPK antibody detected a 72 kilodalton band in wild-type *C. elegans*, which was absent in animals carrying an *aak-2* null mutation (**S1B Fig**), as shown previously [48]. Consistent with the known phosphorylation of AMPK in the setting of high AMP/ATP ratios, ATP levels were low (**S1C Fig**) and phospho-AMPK levels were high (**S1D Fig**) in animals mutant for *pink-1*, the *C. elegans* PARK6 homolog, which is required for mitochondria maintenance [49]. These experiments demonstrate the specificity of the antibody for phosphorylated AAK-2, and the reliability of p-AMPK in reporting the anabolic versus catabolic state of the animal.

Surprisingly, despite the low ATP levels measured during DTS, phosphorylated AMPK was decreased during lethargus/DTS as well as during SIS (**Fig 1B and 1D**). These results suggest that AMPK is not activated in whole animals during these sleep states despite low whole animal ATP levels, suggesting that the animal is in an overall anabolic metabolic state. However, we cannot exclude the possibility that AMPK is activated in specific cells.

Our ATP measurements show that sleep behavior correlates with dropping ATP levels. However, it remained unclear whether sleep behavior causes the ATP drop or whether sleep is a response to ATP drop, i.e. sleep is an attempt by the animal to conserve energy. To distinguish between these possibilities, we developed a pharmacogenetic approach to sleep deprive animals. We expressed a histamine-gated chloride channel in the sleep-promoting RIS neuron [42], and then cultivated animals on histamine during sleep. This approach works because histamine is not used by *C. elegans* as a neurotransmitter [50]. RIS is required for movement quiescence during lethargus/DTS and, in addition, is required for quiescence during UV induced SIS (**S8A-B Fig**). Pharmacogenetic silencing of RIS resulted in a 35.0% ± 10.1 (mean ± SEM, n=6) reduction in DTS (**S1E Fig**) and 72.3% ± 9.5 (mean ± SEM, n=12) reduction in SIS (**Fig 1E, 1G and S1F Fig**). We compared ATP levels during SIS of animals in which sleep was pharmacogenetically reduced to animals expressing the histamine-gated chloride channel but not exposed to histamine. There was no effect of histamine itself on ATP levels. ATP levels were reduced specifically by preventing sleep (**Fig 1G**). These data suggest that the sleep state is not causing the drop in ATP levels; rather, it is a response to increased energetic demands, in an attempt to conserve energy.

When subjected to reduced nutrient intake or to increased nutrient expenditure, animals break down fat stores to release energy for use by all cells. Because during lethargus/DTS, *C. elegans* synthesize and secrete a cuticle [45], an energetically expensive task, we asked whether fat levels during DTS were depleted faster than can be explained by fasting alone. In support of the high energetic demands during lethargus/DTS, we observed a depletion of fat stores. When we fasted awake animals for an hour during the middle of the L4 stage, their fat stores decreased, but animals in lethargus/DTS showed a greater reduction of fat stores (**Fig 2A-B**), suggesting that energetic demands are indeed increased during lethargus/DTS. Thus, a reduction in energy use in the neuromuscular system by sleeping does not fully compensate for the overall energetic demands during DTS and SIS sleep.

**Fig 2.**
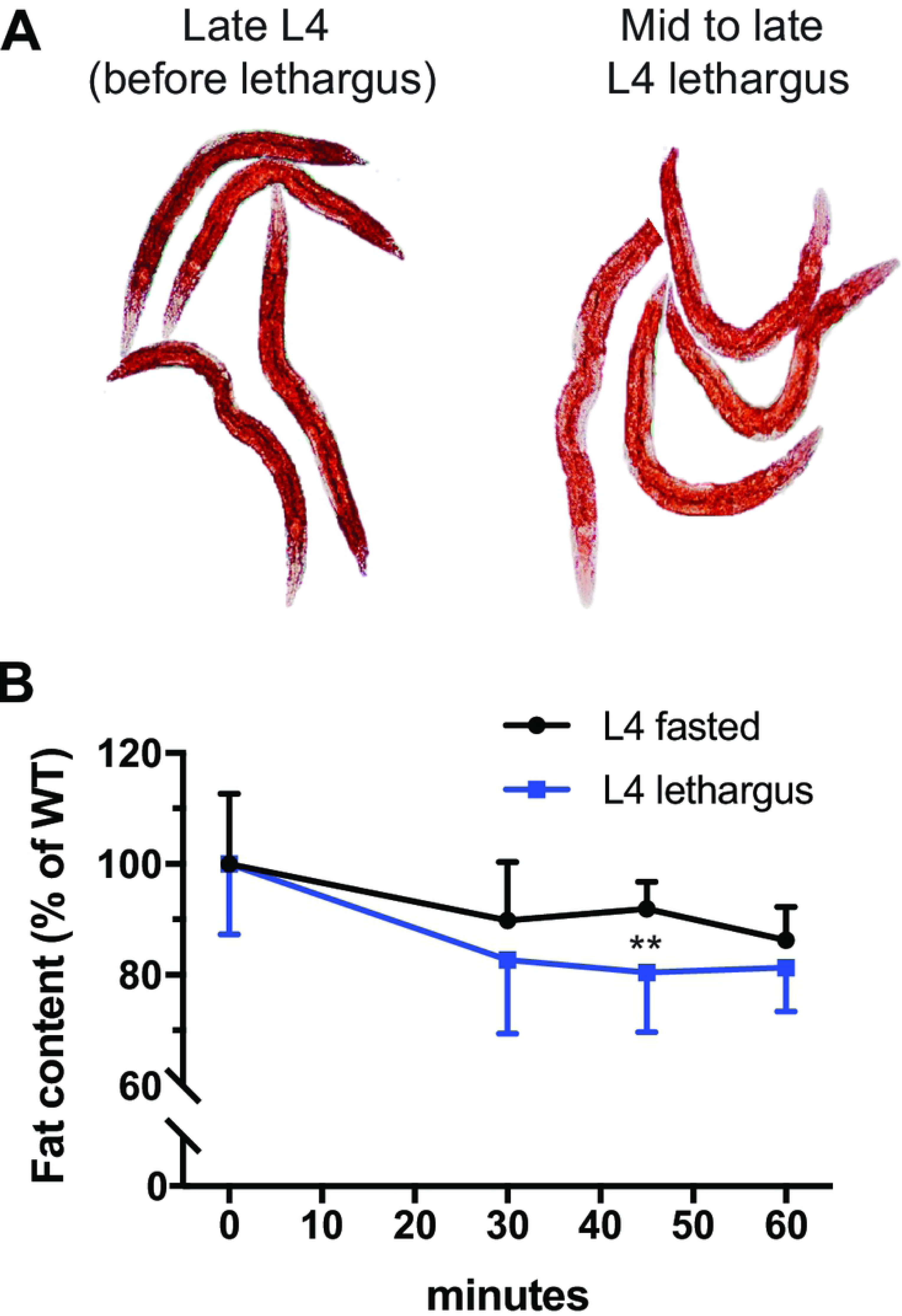
Fat stores are depleted during DTS lethargus sleep. **(A)** Representative images of late L4 larvae taken before lethargus (left image) and L4 larvae during mid to late lethargus (right image). Animals were fixed and stained with Oil Red O, which stains fat droplets. **(B)** Fat content measured with fixative Oil-Red O staining of wild-type L4 larvae during non-lethargus L4 fasting and during lethargus/DTS (n=15 animals per time-point). Fat levels during lethargus/DTS are depleted faster than fat levels of fasting non-lethargus animals. Data are represented as a percentage of non-lethargus fasting animals ± SEM. ** p<0.01, Student’s two-tailed *t* test.

Together, these results indicate that *C. elegans* sleep, both during DTS and during SIS, is associated with a net negative energy balance, where ATP consumption is greater than ATP synthesis and that sleep is an energy conserving state.

### KIN-29 regulates sleep, metabolic stores, and energy charge

We sought to identify molecular signals that mediate this metabolic regulation of sleep. Because SIK3 is a key regulator of sleep [27] and metabolism [34, 51, 52], we reasoned that SIKs may coordinate both metabolism and behavioral state (sleep/wake). In contrast to mammals or *Drosophila*, which have three or two genes respectively encoding SIK proteins, *C. elegans* has only one called KIN-29 **(S2A Fig**) thereby simplifying genetic manipulations.

We assessed sleep phenotypes of *kin-29* null mutants. Consistent with a prior study [27], *kin-29* mutants have reduced lethargus/DTS (**Fig 3A-B and S2B-C Fig**). In addition, *kin-29* mutants had reduced stress-induced sleep (SIS) as determined by movement and feeding quiescence of animals exposed to UV radiation (**Fig 3F-G and S2D Fig**) or to heat stress (**S2E-F Fig**). Together with prior observations of quiescence defects in the setting of satiety [31], these data suggest that *kin-29* is generally required for sleep.

**Fig 3.**
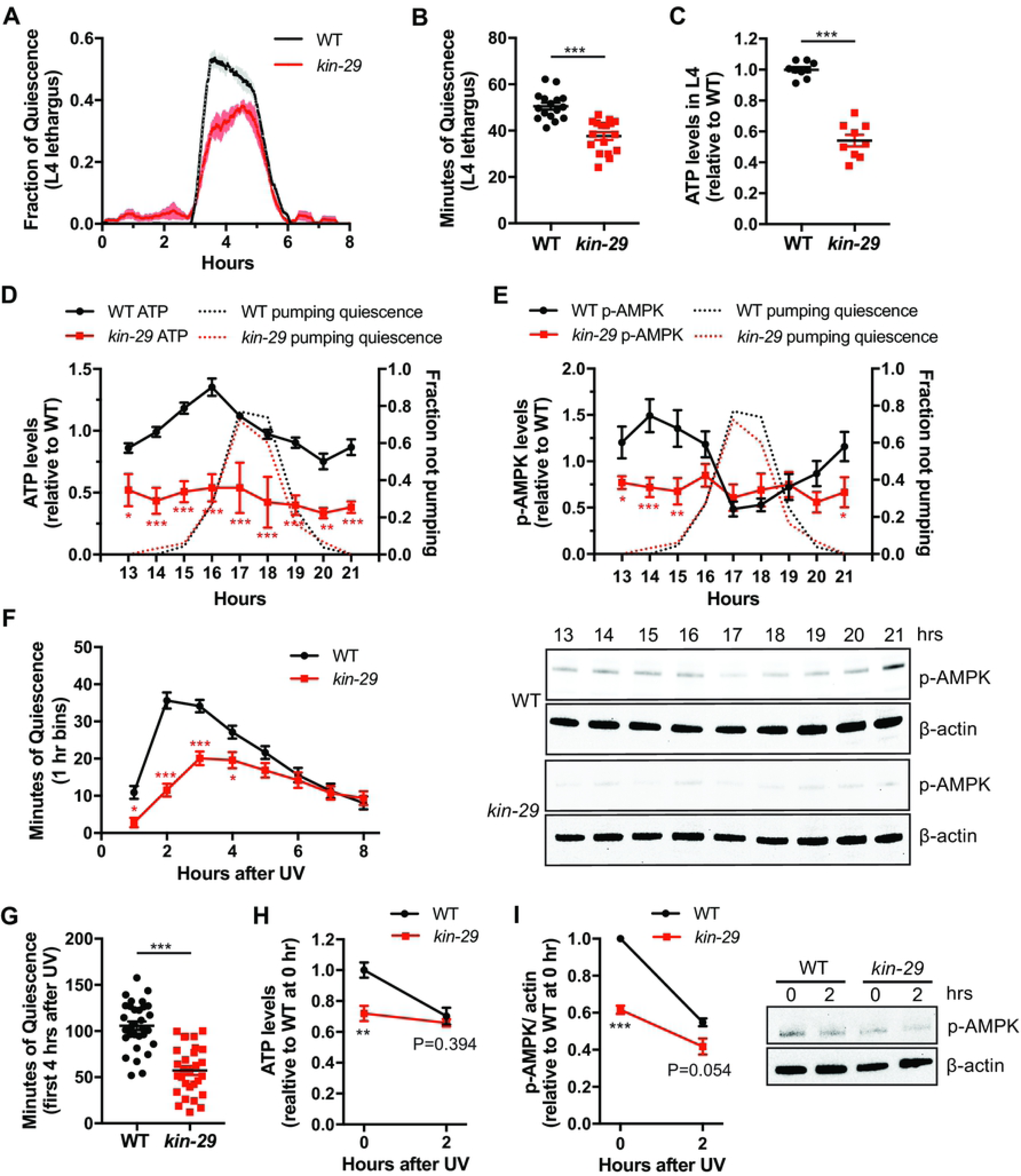
*kin-29* mutants have reduced DTS and SIS, and low ATP and p-AMPK levels. **(A)** Fraction of body movement quiescence of wild-type and *kin-29* null mutant animals. Data are represented as a moving window of the fraction of a 10-min time interval spent quiescent of n=6 animals for each trace. Wild-type and *kin-29* mutants were analyzed on the same WorMotel. The x-axis represents hours from the start of recording in the late fourth (L4) larval stage. **(B)** Total body movement quiescence during L4 lethargus/DTS is lower in *kin-29* null mutants compared to wild-type (n=16-17 animals combined from 3 separate experiments). Data are represented as the mean ± SEM. *** p<0.001, Student’s two-tailed *t* test. **(C)** Levels of total body ATP (nM ATP/μg protein) measured in wild-type mid L4 larvae. Data are normalized to wild-type and represented as the mean ± SEM of 9 experiments. *** p<0.001, Student’s two-tailed *t* test. **(D)** Levels of total body ATP (nM ATP/μg protein) in wild-type and *kin-29* null mutant animals measured before, during and after L1 lethargus/DTS. Data are normalized to the average value of the wild-type time-course. Graphs show the mean ± SEM of 7 experiments for wild-type, and 2 experiments for *kin-29* mutants. Statistical comparisons were performed with a 2-way ANOVA using time and genotype as factors, followed by post-hoc pairwise comparisons at each time point to obtain nominal p values, which were subjected to a Bonferroni correction for multiple comparisons. ***, ** and * indicates corrected p values that are different from wild-type at p<0.001, p<0.01 and p<0.05, respectively. **(E)** Levels of total body p-AMPK (p-AMPK relative to actin) in wild-type and *kin-29* null mutant animals measured before, during and after L1 lethargus/DTS (A). Data are normalized to the average value of the wild-type time-course period. Graphs show the mean ± SEM of 3 experiments for wild-type and *kin-29* mutants. Statistical comparisons were performed by an unpaired multiple comparison *t*-test with Holm-Sidak correction. ***, ** and * indicate corrected p values that are different from *kin-29* mutants at p<0.001, p<0.01, and p<0.05, respectively. Representative Western blots are shown below the graph. **(F)** Time-course of body movement quiescence per hour for wild-type and *kin-29* null mutant animals (n=32 animals for each genotype) after UVC irradiation (1,500 J/m^2^). Graphs show the mean ± SEM. Statistical comparisons were performed with a 2-way ANOVA using time and genotype as factors, followed by post-hoc pairwise comparisons at each time point to obtain nominal p values, which were subjected to a Bonferroni correction for multiple comparisons. *** and * indicates corrected p values that are different from wild-type at p<0.001 and p<0.05, respectively. This is one replicate of an experiment run more than three times. **(G)** Minutes of body movement quiescence during the first 4 hours after UVC exposure/SIS is lower in *kin-29* null mutants compared to wild-type (n=32 animals for each genotype) as determined from the time-course data in F. Data are represented as mean ± SEM. *** p<0.001, Student’s two-tailed *t* test. This is one replicate of an experiment run more than three times. **(H)** Total levels of body ATP levels (relative intensity of nM ATP/μg protein) measured at maximum quiescence (2 hours) after UVC exposure in wild-type animals and *kin-29* null mutants. ATP levels of wild-type and *kin-29* null mutants are not different 2 hours after UVC irradiation. Data were normalized to wild-type without UVC irradiation (0 hours). Graphs show the mean ± SEM of 4-6 experiments. ** p<0.01, Student’s two-tailed *t* test. **(I)** Total p-AMPK levels (relative intensity of p-AMPK/actin) measured at maximum quiescence (2 hours) after UVC exposure in wild-type and *kin-29* null mutants. Graphs show the mean ± SEM of 3 experiments. *** p<0.001, Student’s two-tailed *t* test. A representative Western blot is shown adjacent to the graph.

Based on our above analysis indicating that ATP levels inversely correlate with sleep drive, there are at least two possibilities for the reduced sleep of *kin-29* mutants. First, it is possible that cellular energy levels are high in *kin-29* mutants, thereby reducing sleep drive. Alternatively, it is possible that *kin-29* mutants have low ATP levels and high sleep drive but are defective in the sleep response to low ATP levels. We found that both ATP and p-AMPK levels were lower in *kin-29* mutants than in wild-type controls both during the fourth larval (L4) stage (**Fig 3C)** and during the adult stage (**S2G-H Fig**). Consistent with our whole animal tissue extract ATP determinations, a validated luminescence assay for *in vivo* ATP levels [53] showed reduced ATP levels in *kin-29* adult animals (**S2G Fig**).

Animals with reduced cellular energy, either due to reduced food intake (e.g. *eat-2* mutants [54-56]), or reduced ability to liberate energy from food [57], forage hyperactively in the presence of ample food and leave the bacterial lawn frequently [58, 59]. As predicted by our measurements of low ATP levels, *kin-29* mutants left the bacterial lawn more frequently than wild-type animals (**S3A Fig**). Therefore, both biochemically and behaviorally, *kin-29* mutants show evidence of low cellular ATP levels

These results suggested that in *kin-29* mutants, ATP production is reduced, ATP consumption is increased, or both. We examined the total ATP and p-AMPK levels before, during and after lethargus/DTS. In contrast to the dynamic ATP levels during larval development observed in wild-type animals (**Fig 1A**), both the ATP and p-AMPK levels remained constant and low in *kin-29* mutants across DTS/lethargus (**Fig 3D-E**) and in SIS (**Fig 3H-J**). These results suggest that KIN-29 is required to respond to low cellular energy levels, and that this response may be required to promote sleep.

ATP is generated by break down of macromolecules such as triglycerides [60]. During cultivation of the animals, we noted that *kin-29* mutants had darker intestines than wild-type animals when viewed under bright field stereomicroscopy. An optically dense intestinal phenotype has been reported to correlate with elevated fat stores [61-63]. We therefore hypothesized that *kin-29* mutants had increased fat stores.

To test this hypothesis, we measured fat levels in *kin-29* null mutants using multiple methods including fixative Oil-Red O, fixative Nile-Red staining (O’Rourke, Cell Metabolism, 2009), and measurement of triacylglycerides (TAG) in worm extracts. Using these fat ascertainment methods, we observed increased fat stores in *kin-29* mutants (**Fig 4A-B**). The *kin-29* increased fat phenotype was present throughout the animal life span from the first larval stage through the adult stage (**S3B Fig**). As controls for our fat ascertainment methods, we observed increased fat in animals mutant for the insulin receptor DAF-2 [62] and decreased fat in animals mutant for the gene *eat-2* [63, 64], which is required for food intake [54, 65] (**Fig 4A-B**). To further characterize the excess fat phenotype of *kin-29* mutants, we used a fluorescent *gfp*-reporter that marks the surface of lipid-droplets (DHS-3::GFP) [66]. *kin-29* mutants had increased number and size of lipid droplets in comparison to wild-type animals (**S3C Fig**).

**Fig 4.**
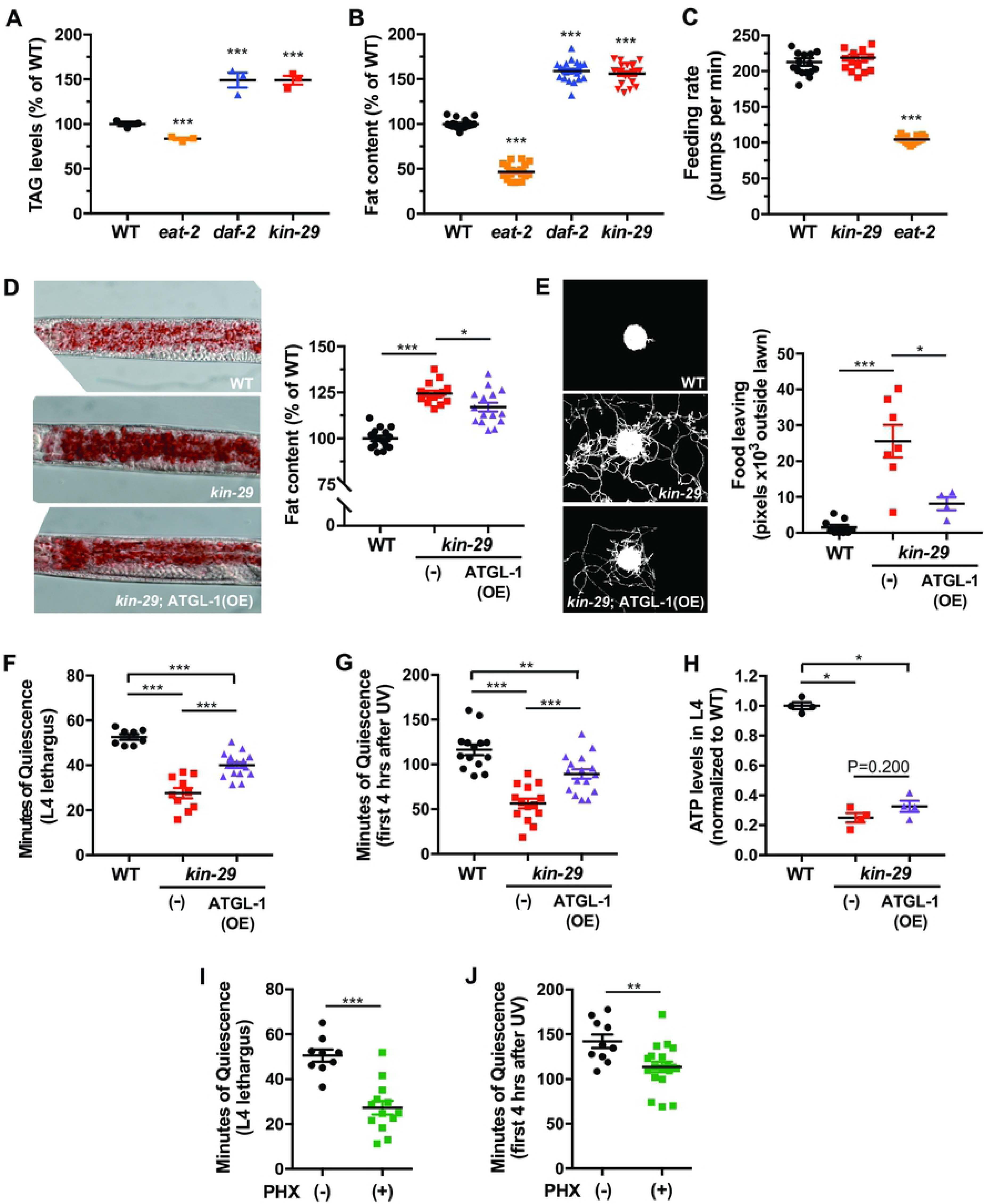
*kin-29* mutants have increased total body fat and food-leaving behavior, which are partially suppressed by ATGL-1 overexpression. **(A)** Total triglyceride levels (μg TAG/μg protein) was quantified for each genotype. The temperature sensitive *daf-2* insulin receptor mutant was raised at 15°C (permissive) and shifted to 20°C (restrictive) at the early L4 larval stage before TAG measurement during the adult stage. Data are represented as the average percentage of total TAG in wild-type controls ± SEM of 3 experiments. *** indicate values that are different from wild-type at p<0.001, Student’s two-tailed *t* test. **(B)** Fat content measured with fixative Oil-Red O staining for each indicated genotype. Data are presented as the average percentage of total body fat in wild-type controls ± SEM (n=19-21 animals). *** indicate values that are different from wild-type at p<0.001, Student’s two-tailed *t* test. **(C)** The feeding rate of *kin-29* null mutants is not different from that of wild-type control animals. Feeding rate was measured as the number of feeding motions (pumps per minute) in the presence of food for each indicated genotype with *eat-2* feeding defective mutants showing as expected reduced feeding rate. Mean ± SEM (n=16 animals). *** indicate values that are different from wild-type at p<0.001, Student’s two-tailed *t* test. **(E)** ATGL-1 overexpression reduces the increased fat stores of *kin-29* null mutants. Fat content was measured with fixative Oil-Red O staining for each indicated genotype, and data are represented as the average percentage of total body fat in wild-type control animals ± SEM (n=15 animals per group). ATGL-1(OE) indicates overexpression of the ATGL-1 adipose triglyceride lipase. Representative images are shown of animals fixed and stained with Oil Red O. * p<0.05, *** p<0.001, Student’s two-tailed *t* test. **(F)** ATGL-1 overexpression (OE) reduces the increased food leaving behavior of *kin-29* null mutants. Food leaving was measured as the area of exploration with each data point representing tracks from a population outside the bacterial lawn. Each data point represents the average number of pixels outside of bacterial lawn of 5-7 animals and the horizontal line represents the mean ± SEM of individual experiments. Representative images are shown of food-leaving behavior, in which frames from a 12-hour video were collapsed into a single image. * p<0.05, *** p<0.001, Student’s two-tailed *t* test. **(G-H)** ATGL-1 overexpression (OE) in the intestine partially restored the reduced L4 lethargus/DTS (G) and UVC-induced quiescence (H) of *kin-29* null mutants. Data are represented as the mean ± SEM with n=8-17 animals for DTS and n=14-16 animals for SIS. *** p<0.001, ** p<0.01, Student’s two-tailed *t* test. **(H)** The reduced ATP levels (nM ATP/μg protein) in L4 larvae of *kin-29* null mutants are not restored by ATGL-1 overexpression. Data are normalized to wild-type and represented as the mean ± SEM of 4 experiments. * p<0.05, Student’s two-tailed *t* test. **(I-J)** Perhexiline treatment results in a reduction in body movement quiescence during L4 lethargus/DTS (I) and after UVC exposure/SIS (J) in wild-type animals. Total minutes of body movement quiescence during L4 lethargus, and during the first 4 hours after UVC exposure in adults in the presence (+) of 1 mM perhexiline (PHX) or in the absence (-) of perhexiline (PHX). Shown is one representative replicate of an experiment run at least four times. Data are represented as mean ± SEM. *** p<0.001, Student’s two-tailed *t* test.

The fat phenotype of *kin-29* mutants is not explained by increased food intake, because feeding behavior as measured by the frequency of pharyngeal contractions (**Fig 4C**) and by the uptake of fluorescence microspheres (**S3D Fig**) [67] was not elevated in *kin-29* mutants when compared to well-fed control wild-type animals.

In summary, these fat assessment methods all show elevated fat stores despite normal food intake in *kin-29* mutants and reduced ATP levels.

### Sleep defects of KIN-29 mutants are corrected by liberation of energy stores

One explanation for the mutant sleep and metabolic phenotypes is that *kin-29* is required to respond to low cellular ATP levels by promoting in parallel both sleep and the liberation of energy from fat stores. An alternative explanation is that fat break down is the signal for sleep and that *kin-29* mutants do not liberate fat energy stores when ATP levels drop. If this second, linear pathway, explanation is correct, then it should be possible to bypass the need for *kin-29* in promoting sleep by using a genetic manipulation that liberates fat directly.

The adipose triglyceride lipase-1 (*atgl-1)* encodes the *C. elegans* orthologue of the rate limiting enzyme in mammalian fat breakdown [68, 69], and is expressed in the *C. elegans* intestinal cells that store fat [70]. We over-expressed ATGL-1 and assessed both cellular energy stores and sleep behavior. We observed a reduction of body fat stores (**Fig 4D**), indicating that the ATGL-1 over-expression achieved the intended goal. ATGL-1 partially restored the defective DTS and SIS sleep phenotype of *kin-29* mutants (**Fig 4F-G and S4A-B Fig**) and corrected the food leaving phenotype (**Fig 4E**), but did not cause an increase in ATP levels (**Fig 4H).** This result suggests that it is the liberation of fat from intestinal cells by ATGL-1 over-expression and not the increase in ATP levels that promotes sleep and reduced food leaving.

We hypothesized that the mechanism by which ATGL-1 over expression promotes sleep is via beta oxidation of the liberated fatty acids. To test this hypothesis, we used the carnitine palmitoyltransferase (CPT) inhibitor perhexiline (PHX) to block fatty acid oxidation [71] (**S5A Fig**). We found that PHX treatment reduced body movement quiescence during lethargus/DTS (**Fig 4I and S5B Fig**) as well as during SIS (**Fig 4J and S5C Fig**). In addition, PHX had a small but significant suppression of feeding quiescence during SIS (**S5D Fig**). The fraction of feeding quiescent animals after UVC exposure was 59.3% ± 1.4 (mean ± SEM, n=27) in the presence of vehicle and 28.6% ± 1.8 (mean ± SEM, n=28) in the presence of PHX.

These results indicate that KIN-29 senses low ATP levels to signal the intestinal cells to liberate and metabolize fatty acids, which then results in signals to sleep-promoting centers by yet unclear mechanisms.

### A sensory neuron basis for KIN-29 SIK in the metabolic regulation of sleep

Similar to broad expression of *sik* genes in mammals [28], *kin-29* is broadly expressed in both neural and non-neural cells in *C. elegans* [29]. Since fat is stored primarily in *C. elegans* intestinal cells [36], we asked whether the excessive fat phenotype of *kin-29* mutants is explained by intestinal action of KIN-29. Expression of *kin-29* under the intestine-specific *ges-1* promoter did not rescue the excess fat, the food-leaving behavior (**Fig 5A-B and S6A Fig**), or sleep defects of *kin-29* mutants, indicating that *kin-29* does not act in the gut to regulate these phenotypes.

**Fig 5.**
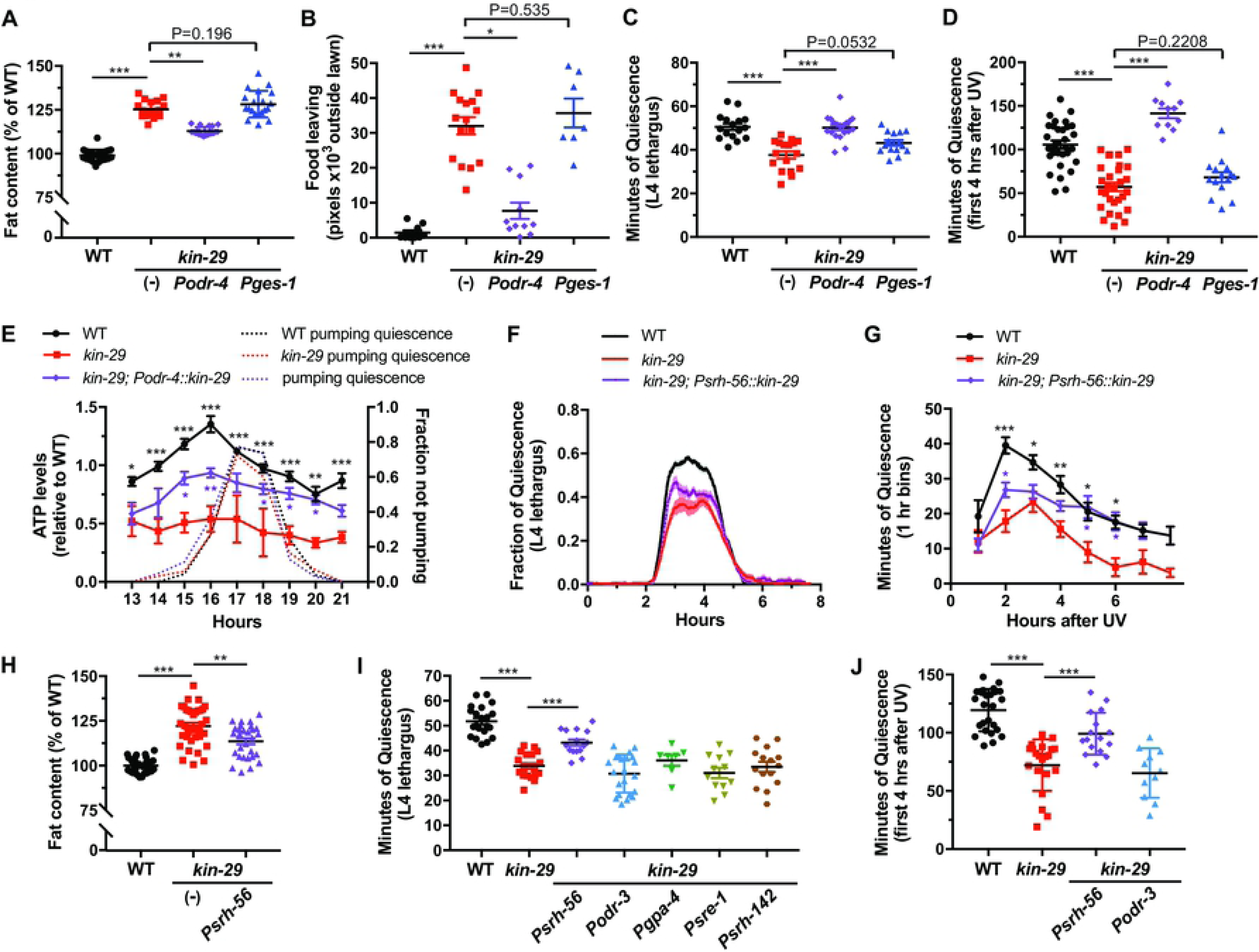
*kin-29* acts in a subset of sensory neurons to regulate DTS and SIS sleep states, fat stores and food-leaving behavior. **(A and B)** Expression of *kin-29* in sensory neurons but not in the gut corrects the increased fat content (I) and increased food-leaving behavioral (J) phenotypes of *kin-29* null mutants. *odr-4* is expressed in 12 pairs of sensory neurons. *ges-1* is expressed in the intestine. Fat content was measured with fixative Nile Red staining, and data are represented as the average percentage of total body fat in wild-type controls ± SEM (n=18-24 animals). Food leaving was quantified as the area of exploration with each data point representing tracks from a population outside the bacterial lawn. Each data point represents the average number of pixels outside of bacterial lawn of 5-7 animals and the horizontal lines represents the mean ± SEM of individual experiments. * p<0.05, ** p<0.01, *** p<0.001, Student’s two-tailed *t* test. **(C and D)** The reduced body movement quiescence of *kin-29* null mutants during L4 lethargus/DTS (C) and after UV exposure/SIS (D) is restored by expressing *kin-29* in *odr-4*-expressing sensory neurons, but not in the *ges-1*-expressing intestinal cells (n>15 animals per group). non-transgenic (-) *kin-29* mutant animals are shown. Data are represented as the mean ± SEM. *** indicate values that are different from wild-type and non-transgenic (-) *kin-29* animals at p<0.001, Student’s two-tailed *t* test. (D) shows the results of one representative experiment run two times. **(E)** Levels of total body ATP (nM ATP/μg protein) measured before, during and after L1 lethargus/DTS of the indicated genotypes. *kin-29* animals expressing *Podr-4::kin-29* restore in part the reduced ATP levels of *kin-29* mutants. *Podr-4::kin-29* is an extrachromosomal transgenic array, and about 20% of siblings of this strain that have lost the array (and are therefore *kin-29* mutants) are included in the ATP measurements, thereby reducing the overall ATP levels. Data are normalized to the average value of the wild-type time-course. Graphs show the mean ± SEM of 2-3 experiments for each genotype. Statistical comparisons were performed with a 2-way ANOVA using time and genotype as factors, followed by post-hoc pairwise comparisons at each time point to obtain nominal p values, which were subjected to a Bonferroni correction for multiple comparisons. ** and * indicate corrected p values that are significantly different between *kin-29* mutants and *kin-29* animals carrying the *Podr-4::kin-29* transgene at p<0.01 and p<0.05, respectively. **(F and G)** Reduced movement quiescence of *kin-29* null mutants during of L4 lethargus/DTS (F) and after UV exposure/SIS (G) are partially restored by expressing *kin-29* in *srh-56*-expressed neurons (ASH, ASJ, ASK). The fraction of quiescence in a 10-minute moving window (F) is shown (n=9 animals for each trace). x-axis represents hours from the start of recording in the late fourth (L4) larval stage. Time-course of body movement quiescence (G) per hour after UVC irradiation (1,500 J/m^2^) with graphs showing the mean ± SEM for each time-point (n=8-10 animals for each genotype). Statistical comparisons were performed by an unpaired multiple comparison *t*-test with Holm-Sidak correction. ***, ** and * indicate corrected p values that are different from *kin-29* mutants at p<0.001, p<0.01, and p<0.05, respectively. **(H)** Fat content measured with fixative Oil-Red O staining. Data are represented as the average percentage of total body fat in wild-type controls ± SEM (n=28-36 animals for each genotype). *** and ** indicate values that are different from wild-type and non-transgenic (-) *kin-29* animals at p<0.001 and p<0.01, respectively, Student’s two-tailed *t* test. **(I and J)** Minutes of body movement quiescence during L4 lethargus/DTS (I) and after UVC exposure/SIS (J) in *kin-29* null mutant animals carrying *kin-29* expression using the indicated promoters compared to wild-type (n>7 animals for each genotype). Data are represented as the mean ± SEM. *** p<0.001, Student’s two-tailed *t* test.

We next assessed a role for *kin-29* in the nervous system. The sensory nervous system of *C. elegans*, similar to the mammalian hypothalamus, plays an important role in sensing nutrient availability and signaling to regulate animal metabolism [72]. Two *kin-29* phenotypes, a small body size and the propensity to enter the dauer diapause stage, are corrected by using the *odr-4* promoter to express the *kin-29* cDNA in 12 pairs of sensory neurons [29]. We tested the hypothesis that the fat storage phenotype too is controlled by *kin-29* acting in these *odr-4*(+) sensory neurons. *odr-4* promoter driven *kin-29* rescued the high fat stores (**Fig 5A and S6A Fig**), lipid droplet morphology (**S3C Fig**) and reduced ATP level phenotypes (**Fig 5E and S6B Fig**) of *kin-29* mutants. In addition, *Podr-4::kin-29*, but not *Pges-1::kin-29*, rescued the defective DTS and SIS sleep phenotypes (**Fig 5C-D and S6C Fig**), and the food-leaving starvation behavior (**Fig 5B**) of *kin-29* mutants.

We next examined the role of KIN-29 function in DTS and SIS in subsets of the 12 sensory neurons defined by the *odr-4* promoter activity (**S7A Fig**). Reconstitution of *kin-29* function in the ASH, ASK, and ASJ sensory neuron pairs using the *srh-56* promoter (**S7B Fig**) partially corrected the DTS and SIS phenotype (**Fig 5F-G**) as well as the fat phenotypes (**Fig 5H and S7C Fig**) of *kin-29* mutants. In contrast, *kin-29* expressed in ASH, AWA, AWB, AWC, and ADF under the control of the *odr-3* promoter, ASI under the control of the *gpa-4* promoter, ADL under the control of the *sre-1* promoter, or ADF under the control of the *srh-142* promoter sensory neuron pairs, did not rescue the sleep phenotypes (**Fig 5I-J and S7D-E Fig**). These data suggest that KIN-29 function in ASK and/or ASJ is important for sleep and lipid homeostasis.

Because the rescue of these phenotypes using the *srh-56* promoter to drive *kin-29* expression is weaker than the rescue using the *odr-4* promoter, *kin-29* likely also functions in other, as yet undefined, *odr-4(+)* neurons to regulate sleep and lipid homeostasis.

Taking together, these data further support a role for *kin-29* acting in sensory neurons to regulate intestinal fat and organismal sleep. Importantly, our data suggest that *kin-29* acts in the same neurons to regulate both fat and sleep, as would be predicted by a linear model in which fat liberation promotes sleep.

### KIN-29 SIK acts upstream of ALA and RIS activation to promote sleep

The *odr-4* gene is not expressed [73] in the two best-characterized interneurons regulating sleep, the ALA [39] and RIS [42] neurons, suggesting that *kin-29* does not act in these sleep executing neurons but rather acts either at a step prior to activation of these neurons, or at a step after activation of these neurons.

Epidermal growth factor (EGF) activates the ALA neuron [74], which regulates SIS by releasing a cocktail of neuropeptides including those encoded by the gene *flp-13* [44, 75]. To determine whether *kin-29* functions upstream or down stream of the ALA neuron, we asked whether the quiescence-inducing effect of EGF activation of ALA is attenuated in *kin-29* mutants. We observed no effect of a *kin-29* null mutation on the quiescence induced by overexpressing EGF (**Fig 6B-C**), supporting the notion that *kin-29* acts upstream of ALA activation by EGF. As expected for a gene acting upstream of ALA activation, the *kin-29* null mutation also did not attenuate the quiescence induced by overexpressing FLP-13 peptides **(S8A-B Fig)**.

**Fig 6.**
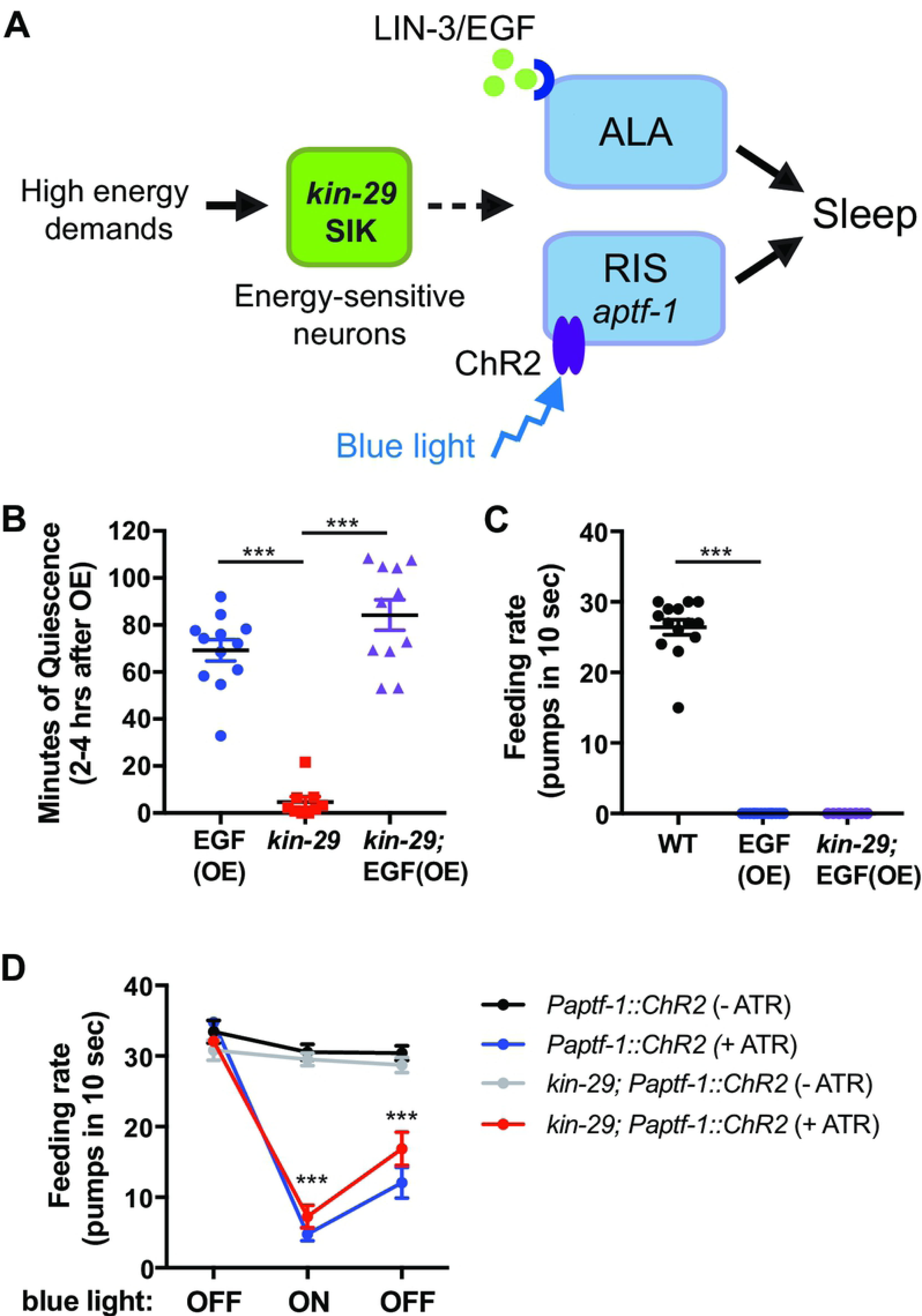
*kin-29* acts upstream of ALA and RIS neurons to regulate sleep. **(A)** Model by which *kin-29*/SIK functions in energy-sensitive sensory neurons upstream of the sleep-promoting RIS and ALA neurons. ALA activation by the LIN-3/EGF or RIS activation by channelrhodopsin2 (ChR2) bypasses the requirement for KIN-29 function in sleep. RIS is also required for movement quiescence during SIS (see **S8C-F Fig). (B and C)** *kin-29* null mutation does not affect the body movement quiescence (A) and reduced feeding rate (B) observed in response to EGF overexpression (OE) (n>9 animals). To induce EGF overexpression, adult animals expressing a *Phsp-16.2::LIN-3C* transgene were heat shocked for 30 min 2 hours prior to analysis of behavior (see **Material and Methods**). Data are represented as mean ± SEM. *** p<0.001, Student’s two-tailed *t* test. B and C are data from one replicate of an experiment run at least two times. **(D)** Optogenetic stimulation of the RIS neuron causes a reduction in feeding rate, which is not dependent on *kin-29*. Wild-type and *kin-29* null mutants expressing *Paptf-1::ChR2* grown either in the presence or absence of all-*trans* retinal (ATR) were exposed to blue light (ON) (see **Material and Methods**). Pumps were counted during a 10-second window, before, during, and after exposure of transgenic animals (n=9-20) to blue light. Data are represented as the mean ± SEM for each condition. *** p<0.001, Student’s two-tailed *t* test.

DTS is primarily controlled by the RIS neuron, which releases neuropeptides encoded by the gene *flp-11* [76]. In addition to the requirement of ALA for SIS, we observed that RIS is required for body movement quiescence and, to a lesser extent, for feeding quiescence during SIS **(Fig 6A and S8C-F Fig)**. To ask if KIN-29 functions upstream of RIS, we crossed *kin-29* mutants into a strain expressing channelrhodopsin2 (ChR2) under the *aptf-1* promoter to activate RIS [42]. Illuminating adult worms expressing *Paptf-1::ChR2* with blue light leads to cessation of pumping when worms are treated with the ChR2 cofactor *all-trans retinal* (ATR), but no change in pumping rate in non-ATR controls [42]. The *kin-29* null mutation did not impair ATR-dependent reduction in pumping in response to optogenetic activation of *aptf-1* expressing neurons (**Fig 6D)**, indicating that KIN-29 acts upstream of RIS.

Together, these results indicate that KIN-29 functions in energy-sensitive sensory neurons upstream of the sleep-promoting ALA and RIS neurons. These data are again consistent with a linear model in which *kin-29*, in response to dropping ATP levels, promotes fat liberation, which in turn promotes sleep via activation of ALA and/or RIS.

### KIN-29 SIK acts in sensory neuron nuclei to regulate sleep

Under standard growth conditions, KIN-29 localizes to the cytosol, but in response to cell stress induced by high heat exposure, KIN-29 moves into the nucleus [29]. It regulates gene transcription via interaction with the nuclear factors MEF-2 and HDA-4 [77]. In contrast, the mammalian KIN-29 homolog SIK3 protein has been proposed to act in the cytosol to phosphorylate synaptic proteins [30]. To determine where KIN-29 acts to regulate sleep, we began by assessing its subcellular localization during sleep.

We found that one hour prior to L1 lethargus as well as one hour after L1 lethargus, KIN-29 expressed in *odr-4(+)* neurons is strictly cytoplasmic (**Fig 7A-B**). By contrast, during early, mid, and late L1 lethargus, KIN-29 localizes to the nucleus of a subset of *odr-4(+)* neurons (**Fig 7A-B**). These data lead us to hypothesize that *kin-29* functions in the nucleus to regulate sleep. Based on this hypothesis, we would predict that a *kin-29* mutant that fails to translocate to the nucleus would have a defective regulation of sleep. We were able to test this prediction by studying the function of a KIN-29 protein with a conserved Serine 517 mutated to Alanine (**S9A Fig**). The motivation for making this particular mutant was the observation that a homologous change in the mouse SIK3 gene results in a sleepy phenotype [78]. While we did not observe a sleepy phenotype in the *kin-29(S517A)* mutants, we found that KIN-29(S517A) mutant protein did not move to the nucleus during lethargus (**Fig 7A-B**); moreover, it did not rescue the sleeping-defective of *kin-29* null mutants (**Fig 7C**). KIN-29(S517A) stayed in the cytosol even after heat-shock (**S9B Fig**), which strongly promotes nuclear localization of wild-type KIN-29 [29]. KIN-29(S517A) was otherwise functional because it rescued the small body size phenotype of *kin-29* mutants (**S9C Fig**).

**Fig 7.**
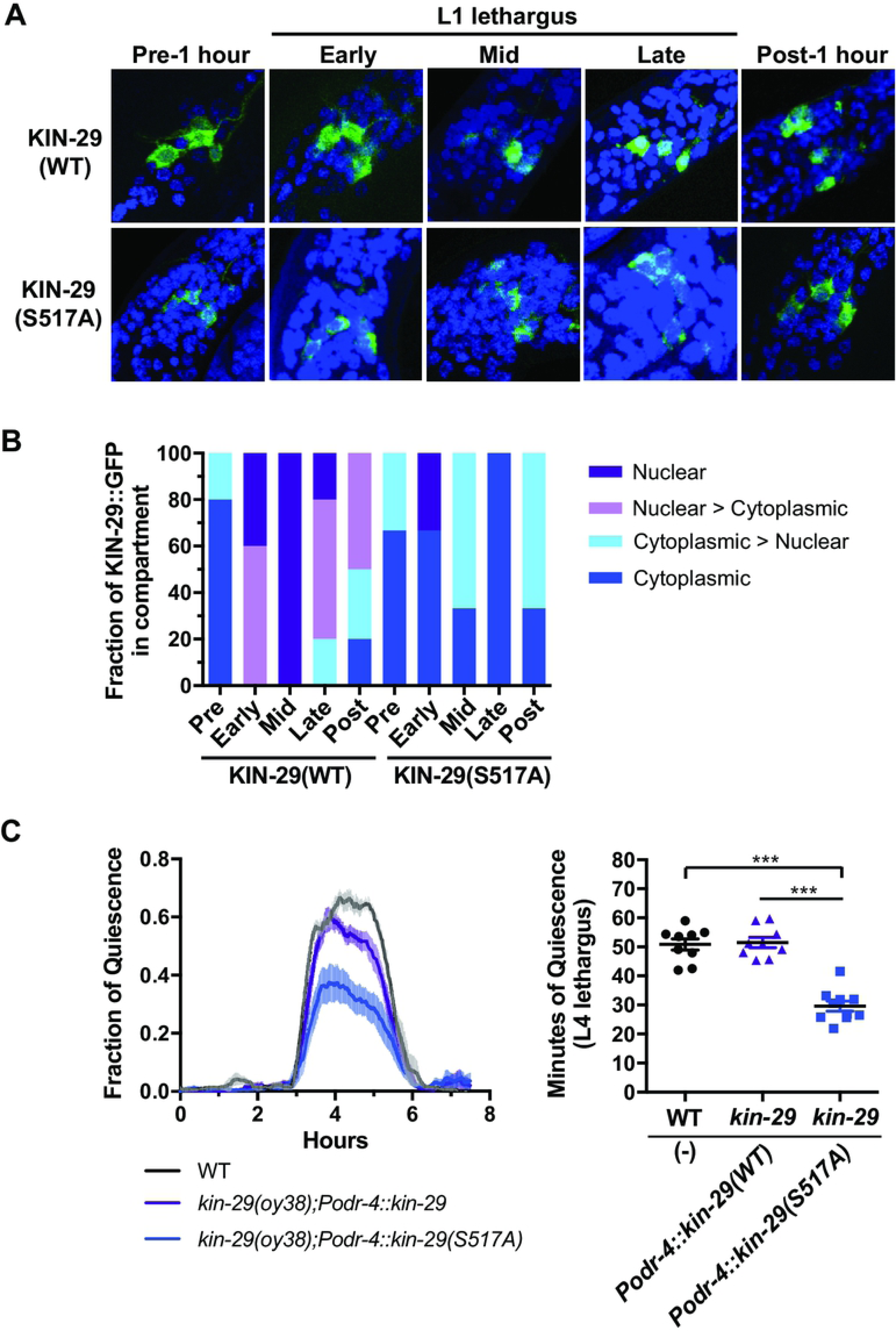
KIN-29(S517A) mutant does not translocate to the nucleus during sleep and does not rescue the short sleeping phenotype of *kin-29*. **(A)** Confocal images of KIN-29::GFP in *odr-4(+)* neurons (green) of wild-type animals. Nuclei are identified by DAPI staining (blue). One hour prior to L1 lethargus and one hour after L1 lethargus (n=3-5 animals for each condition), KIN-29::GFP is mostly cytoplasmic. During early, mid and late L1 lethargus (n= 5-7 animals for each condition), KIN-29, but not the KIN-29(S517A) mutant, localizes to the nucleus of a subset of *odr-4(+)* neurons. **(B)** Fraction of KIN-29::GFP in the subcellular compartment of *odr-4(+)* neurons by showing the percentage of animals that show fully nuclear, intermediate (nuclear > cytoplasmic, and cytoplasmic > nuclear), and fully cytoplasmic location of GFP. **(C)** The KIN-29(S517A) mutant does not rescue the reduced L4 lethargus/DTS of *kin-29* null mutants. Left graph: The fraction of quiescence in a 10-minute moving window is shown with n=4 animals for the wild-type trace, n=8 animals for the *Podr-4::kin-29(S517A)* trace, and n=6 animals for the *Podr-4::kin-29(WT)* trace. x-axis represents hours from the start of recording in the late fourth (L4) larval stage. Data are represented as the mean ± SEM. Right graph: Total body movement quiescence during L4 lethargus/DTS determined from the time-course data. Data are represented as the mean ± SEM. *** p<0.001, Student’s two-tailed *t* test.

If KIN-29 were indeed acting in the nucleus to regulate sleep, then we would predict that it would genetically interact with nuclear factors. To test this prediction, we tested for genetic interactions between *kin-29* and the class II histone deacetylase HDA-4, which KIN-29 has been shown to phosphorylate and inhibit to regulate gene expression in sensory neurons [77]. Moreover, HDA-4 is found in the nuclei of most cells (van der Linden et al., 2008). To determine whether HDA-4 is also required for the KIN-29 regulation of sleep and fat stores, we studied the phenotype of animals’ mutant for both *kin-29* and *hda-4*. Loss-of-function mutations in *hda-4* corrected the DTS and SIS sleep phenotypes (**Fig 8A-B and S10A-B Fig**), and food-leaving behavior of *kin-29* mutants (**Fig 8C and S10C Fig**), suggesting that *hda-4* is negatively regulated by KIN-29 and acts downstream of *kin-29* to regulate sleep and starvation behavior. Expression of *hda-4* under the control of its own promoter in *kin-29 hda-4* double mutants fully restored the defective sleep phenotype of *kin-29* mutants. Expression of *hda-4* under the control of the *odr-4* promoter partially restored the defective SIS sleep phenotype and fully restored the DTS phenotype of *kin-29* single mutants (**Fig 8A-B and S10B Fig**). These data are consistent with KIN-29 acting on HDA-4 in *odr-4(+)* sensory neurons, but suggest that *hda-4* may have additional roles elsewhere in the animal.

**Fig 8.**
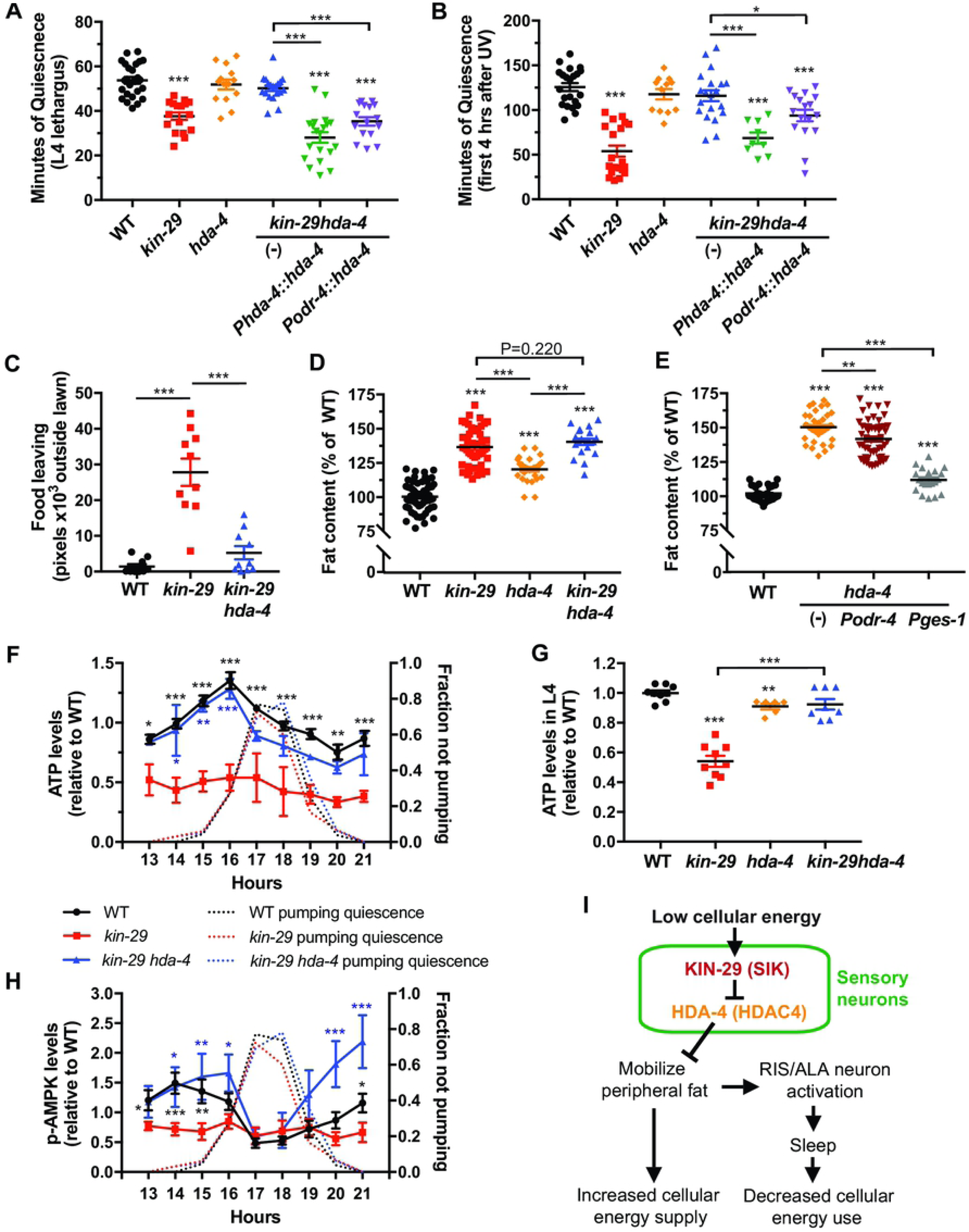
*hda-4* acts downstream of *kin-29* in sensory neurons to control the metabolic regulation of sleep. **(A and B)** An *hda-4* null mutation corrects *kin-29* null mutant sleep to wild-type levels. Restoring *hda-4* in *kin-29hda-4* double mutants under control of the *hda-4* promoter (*Phda-4::hda-4*) or the *odr-4* promoter (*Podr-4::hda-4*) results in reemergence of the *kin-29* quiescence defects during DTS (A) (n=17-28 animals) and SIS (B) (n=9-23 animals). Data are represented as the mean ± SEM. *** and * indicate values that are different from wild-type and non-transgenic (-) *kin-29 hda-4* double mutant animals at p<0.001 and p<0.05, respectively, Student’s two-tailed *t* test. **(C and D)** Fat content and food-leaving behavior for each indicated genotype. (C) Food leaving was quantified as the area of exploration with each data point representing tracks from a population outside the bacterial lawn. Each data point represents the average number of pixels outside of bacterial lawn of 5-7 animals and the horizontal line represents the mean ± SEM of individual experiments. (D) Fat content was measured with fixative Oil-Red O staining, and data are represented as the average percentage of total body fat in wild-type controls ± SEM (n=20-63 animals). *** indicate values that are different from wild-type at p<0.001, Student’s two-tailed *t* test. **(E)** Fat content measured with fixative Nile Red staining for each indicated genotype (n=19-52 animals). Data are represented as the average percentage of total body fat in wild-type controls ± SEM. *** and ** indicate values that are different from wild-type and non-transgenic (-) *hda-4* animals at p<0.001 and p<0.01, respectively. **(F)** Levels of total body ATP (nM ATP/μg protein) in wild-type, *kin-29* single mutants, and *kin-29 hda-4* double mutant animals measured before, during and after L1 lethargus/DTS. Data are normalized to the average value of the wild-type time-course. Graphs show the mean ± SEM of 2-7 experiments for ATP. Statistical comparisons were performed with a 2-way ANOVA using time and genotype as factors, followed by post-hoc pairwise comparisons at each time point to obtain nominal p values, which were subjected to a Bonferroni correction for multiple comparisons. ***, ** and * indicates corrected p values that are different from *kin-29* mutants at p<0.001, p<0.01 and p<0.05, respectively. **(G)** Mutations in *hda-4* restore the reduced ATP levels (nM ATP/μg protein) in L4 larvae of *kin-29* null mutants. Data are normalized to wild-type and represented as the mean ± SEM of 6-9 experiments. *** p<0.001, ** p<0.01, Student’s two-tailed *t* test. **(H)** Levels of total body phosphorylated AMPK (relative intensity of p-AMPK/actin) in wild-type, *kin-29* single and *kin-29hda-4* double mutant animals measured before, during and after L1 lethargus/DTS. Colors denoting each genotype are the same as those used in panel F. Data are normalized to the average value of the wild-type time-course. Graphs show the mean ± SEM of 3-6 experiments for p-AMPK. Statistical comparisons were performed by an unpaired multiple comparison *t*-test with Holm-Sidak correction. ***, ** and * indicate corrected p values that are different from *kin-29* mutants at p<0.001, p<0.01, and p<0.05, respectively. **(I)** Proposed mechanism of metabolic sleep regulation. KIN-29 SIK acts to respond to low energetic stores by signaling to non-neural cells to liberate fat, which in turn promote sleep behavior.

Although the *hda-4* mutation did not suppress the elevated fat phenotype of *kin-29* mutants (**Fig 8D and S10C Fig**) the interpretation of this negative result is confounded by the observation that animals that were mutant only for *hda-4* had increased body fat (**Fig 8D and S10C Fig**), and the increased fat phenotype of *hda-4* single mutants was nearly fully corrected by expressing *hda-4* in the intestine (**Fig 8E**), suggesting that *hda-4* functions in the intestine to control fat stores. We also observed a small but statistically significant reduction in the fat phenotype of *hda-4* mutants when we expressed *hda-4* in *odr-4(+)* neurons (**Fig 8E**), suggesting that *hda-4* functions also in neurons to regulate body fat stores. Mutation in *hda-4* corrected both the low ATP (**Fig 8F-G and S10D Fig**) and p-AMPK abnormalities (**Fig 8H and S10D-E Fig**) of *kin-29* mutants.

Collectively, the genetic interactions between *kin-29* and *hda-4* and the subcellular distribution of KIN-29 during DTS support the notion that KIN-29 acts in the nucleus to regulate sleep.

## DISCUSSION

While the focus of much of sleep function and regulation research has been on brain neurons [79], extensive observations, both basic [80] and clinical [81, 82], demonstrate a role for metabolic sleep regulators outside the nervous system. Metabolic advantages of sleep include conservation of energy [15, 83], proper allocation of metabolic resources [18], temporal segregation of incompatible cellular activities [84], and energetic efficiency [80]. The observation of *C. elegans* sleep in the setting of starvation [4, 32] support a role for sleep in energy conservation, and the observation of sleep following cell injury [39] and during lethargus [38], when nervous system activity is dampened [10-12] support a role for sleep in the reallocation of metabolic resources from excitable cell function to anabolic and repair functions outside the nervous system. In support of an energy conserving role for sleep, we found that preventing sleep results in a drop in ATP levels (**Fig 1G**).

Central nervous system neurons control sleep in a top-down fashion [85, 86], but bottom-up metabolic signals from glia [17, 24, 87, 88], muscle cells [32, 88-90] and adipocytes [91, 92] affect activity of sleep regulating neurons. While several gene products have been reported to regulate both metabolism and sleep [4, 9, 21-23, 25, 26, 32, 41, 89, 90, 93, 94], the mechanism of the metabolic regulation of sleep has heretofore remained opaque.

Our data suggests a model (**Fig 8I**) in which low cellular energy charge of the animal is interpreted by the protein kinase KIN-29 SIK. While numerous potential SIK3 substrates in mouse brains were recently identified [30], our genetic data suggests that KIN-29 SIK acts primarily via a single nuclear protein substrate, the type IIa histone deacetylase HDA-4, to regulate sleep. We propose that KIN-29 SIK phosphorylates and inhibits HDA-4 in a small set of neuroendocrine cells. Inhibition of HDA-4 results in de-repression of genes, which in turn results in humoral signaling from neuroendocrine cells to adipocytes to release energy stored as triglycerides. Liberated energy stores then signal to the sleep-promoting neurons ALA and RIS, which trigger organismal sleep. The mechanism of this signaling remains unknown but one possibility is that an increase of energy stores in intestinal cells leads to the release of one or more of intestinal insulins, which then act on the DAF-2 insulin receptor. Supporting such a mechanism are recent reports that signaling by the DAF-2 insulin receptor and the FOXO transcription factor DAF-16 play a role in the promotion of sleep under certain conditions [4, 32, 90]. An alternative possibility is that liberated free fatty acids or their metabolites play a signaling role in regulating sleep, a mechanism that would be similar to sleep regulation by arachidonic acid metabolites in mammals [95]. Finally, a third possibility is that fatty acid catabolism biproducts such as reactive oxygen species promote sleep, as has recently been demonstrated in *Drosophila* [96, 97].

Our finding that the roles of KIN-29 in both fat mobilization and sleep regulation map to a small number of sensory neurons supports the view that fat homeostasis and sleep are mechanistically linked. Further supporting this notion is our observation that a genetic manipulation in gut/adipocyte cells to liberate energy stored as triglycerides promotes sleep in *kin-29* mutant animals.

Our transgenic rescue experiments implicate in the metabolic regulation of sleep by *kin-29* the sensory neuron types ASJ and ASK as well as in one or more of nine other sensory neuron types expressing the gene *odr-4*. While sleep is associated with reduced ATP levels in whole animals, and *kin-29* controls sleep by action in sensory neurons, we do not yet know whether low energy levels are sensed specifically in these sensory neurons or elsewhere in the animal. We favor the possibility that low energy is detected specifically in sensory neurons. Since information processing during wake entails a high energetic cost [98], we speculate that sensory neurons are particularly sensitive to metabolic needs of the animal and thus play a key role in both sensing and reporting on low cellular energy levels. By reducing their activity, sensory neurons then gate sensory information during sleep [10-12].

During times of acute metabolic stress, AMPK activation plays a key role in suppressing energetically expensive anabolic processes and enhancing energy-generating catabolic processes to maintain or restore ATP intracellular levels [99]. Surprisingly, we find that the drop in ATP levels during sleep occurs without activation of AMPK by phosphorylation until after sleep. This finding suggests that turning on catabolic processes through AMPK activation may be maladaptive to the completion of the anabolic process engaged by the animal. That is, the adaptive process of AMPK activation in response to metabolic stress may become maladaptive and harmful under conditions of extreme stress, as previously proposed [100]. We also observed constitutively low levels of phospho-AMPK in *kin-29* mutants. Our observation of low pAMPK levels in *kin-29* loss-of-function mutants are consistent with a recent observation of elevated phospho-AMPK levels in mice harboring a gain-of-function SIK3 mutant [30].

Our phenotypic characterization indicates that, like mouse and *Drosophila* SIK3, KIN-29 is required for sleep. Moreover, like *Drosophila* dSIK [33], KIN-29 is required in neurons to mobilize fat stores from adipocytes. Because KIN-29 is ancestral to all *Drosophila* and mice SIK proteins, it may alone serve functions that are served separately by dSIK and SIK3 in *Drosophila* and by SIK1, SIK2, and SIK3 in mammals.

SIK3 genetic variants are associated with obesity [101, 102]. It would be of interest to know if those obese individuals also have short sleep, as would be predicted by epidemiological studies showing short sleep to be associated with obesity [3, 103]. Within the framework of the linear model we propose for sleep regulation by fat, we suggest that the association between short sleep and elevated fat stores in humans could be explained by chronic obesity promoting short sleep rather than vice versa.

## MATERIAL AND METHODS

### Strains, general animal cultivation and genetic controls

Worms were cultivated on the surface of NGM agar. Unless otherwise specified, worms were fed the *Escherichia coli* strain OP50 [104] or its derivative DA837 [105] and grown in 20°C incubators. All experiments reported here were performed on hermaphrodites. The wild-type strain used was N2, variety Bristol [104]. A complete strain list used in this study is provided in **S1 Table.** Double mutant animals were constructed using standard genetic methods [106], and genotypes were confirmed by genetic linkage (for example, using balancer chromosomes marked with fluorescence), by phenotype, by PCR (for example, identifying small deletions) or by sequencing of a PCR product (for example, identifying single nucleotide changes).

### Generation of plasmids and transgenic animals expressing *kin-29* and *hda-4*

To generate transgenic worms expressing *kin-29* cDNA in different tissues and cells, the coding region of *kin-29* fused at its C-terminus to GFP coding region and the *unc-54* 3’UTR sequence were cloned in the multiple cloning site (MCS) of the pMC70 plasmid (a gift from the Sengupta lab) resulting in the plasmid pSL165 (*kin-29 cDNA::GFP::unc-54 3’UTR*). Next, promoter sequences of either *ges-1* (2.0 kb), *odr-4* (3.1 kb), *odr-3* (1.7 kb), *srh-56* (1.5 kb), *gpa-4* (3.0 kb), *sre-1* (1.5 kb) or *srh-142* (2.0 kb) were cloned at the 5’ end of the *kin-29* cDNA using the 5’ MCS of pSL165.

To generate transgenic worms expressing the *hda-4* gene under control of the *odr-4* and *ges-1* promoters, the *hda-4* cDNA was fused at its C-terminus to the GFP coding region and the *unc-54* 3’UTR sequence and inserted into the second MCS of the pMC70 plasmid. Promoters of *odr-4* (3.2 kb), or of *ges-1* (2.0 kb) were then cloned at the 5’-end of the start ATG of *hda-4*. Oligonucleotides used and generated constructs are listed in **S2 and S3 Table**, respectively. Constructs were injected into N2 worms at a concentration of 20-50 ng/μl along with P*unc-121::RFP* (AddGene) at a concentration of 100 ng/μl as a transgenesis marker to bring the final concentration up to 150 ng/μl. Generated transgenic lines can be found in **S1 Table.**

### Assessment of movement quiescence

Movement quiescence was measured using the 48-well (6×8) WorMotel [107] for stress-induced sleep (SIS) assessments and a 24-well (4×6) WorMotel for developmentally timed sleep (DTS) assays. For DTS experiments, early to mid-fourth larval stage (L4) animals were imaged for 12-18 hours. For SIS experiments, first day-old adult worms were imaged for 8 hours after stress induction by UV or for 2 hours after stress induction by heat shock. Briefly, worms were placed individually onto the NGM agar surface of WorMotel wells together with a thin layer of bacteria. The worms were imaged under dark field illumination provided by a red LED strip. Images were captured every 10 seconds for the duration of recording using ∼8.5 μm/pixel spatial resolution. Images were analyzed by a pixel subtraction [38] using custom Matlab software (https://github.com/cfangyen/wormotel). Movement quiescence was defined as a lack of changed pixels between successive frames.

### Feeding assessment

Feeding was assessed by counting the number of movements (pumps) of the pharyngeal grinder, a tooth-like structure located in the terminal bulb of the pharynx, over the course of a 10 second window under direct observations under 40x-115x magnification of a stereomicroscope. The experimentor was blinded to the genotype/condition of the worm. One pharyngeal pump was defined as a backward movement of the grinder. Animals were considered feeding quiescent if there were no pharyngeal pumps in the 10 second window. In heat stress experiments, feeding was measured at times 0, 15, 30, 45, and 60 minutes after heat shock, and movement quiescence was measured continuously for two hours after heat shock in a separate cohort of worms. In UV stress experiments, feeding was measured 2 hours after UV and movement quiescence was measured continuously using the WorMotel device for 8 hours after UV exposure.

For the assay of microsphere accumulation in the absence of food, worms were exposed to fluorescent polystyrene microspheres of 1.0 μm diameter (Polysciences) as described [67]. In brief, a 100 μl microsphere suspension was mixed with 100 μl S-basel buffer, spread on a 3 cm NMG-agar plate, and left at room temperature for ∼60 min for liquid absorption. An age-synchronized population was grown until the adult stage. First day-old adult worms were washed three times in S-basal buffer, transferred to the microsphere plates, and incubated ∼15 min for uptake of the microspheres. After incubation, worms were quickly washed with M9 buffer to remove excess microspheres, mounted on 2% agar pads containing the anesthetic Na-azide (NaN_3_), and imaged on a Leica DM5500 Nomarski microscope equipped with a Hamamatsu Orca II camera. The fluorescence intensity of microspheres accumulated in the worm gut was quantified using Volocity software (PerkinElmer)

### SIS induction by UV irradiation and heat shock

UV induced sleep assays [43] were performed by exposing first day-old adult worms to 1500 J/m^2^ UVC irradiation (254 nm) in a Spectrolinker™ XL-1500 (Spectroline). For the UV exposure, the worms were housed either in a WorMotel chip placed in an uncovered plastic 10 cm Petri dish or on the agar surface of an uncovered 5.5 cm Petri dish filled with NGM agar. A thin layer of *E. coli* DA837 or OP50 bacteria was spread onto the surface of the NGM agar immediately before the experiment in order to minimize growth of the bacteria, which could act to screen the worms from the UV. Heat shock induced sleep assays were performed by submerging first day-old day adult animals in a circulating water bath to 35°C for 30 minutes. During the submersion, the worms were housed on the agar surface of a 5.5 cm diameter Petri dish containing 11 mL NGM agar or of a WorMotel placed in an empty plastic 10 cm Petri dish sealed with Parafilm.

### Induction of EGF and FLP-13 overexpression

To induce expression of LIN-3C(EGF) or FLP-13 peptides, first day old adult animals carrying *Phsp-16.2::lin-3C or Phsp-16.2::flp-13* transgenes were housed on the agar surface of a 5.5 cm diameter agar surface (and 11 mL volume of NGM agar) or of a WorMotel and submerged in a circulating water bath at 33°C for 30 minutes. Feeding quiescence was measured 2-2.5 hours after heat-induced transgene induction whereas movement quiescence was measured continuously on the WorMotel device for 8 hours after heat exposure.

### Assessment of total body fat stores

Oil-Red O fixative staining was performed as described [108]. Briefly, well-fed worms were age-synchronized by the bleaching method and grown at 20°C on NGM plates seeded with *E. coli* OP50. L4-staged worms were collected with dH_2_O and washed over a 15 μm nylon mesh filter to remove any bacteria. Worms were transferred to 1.5 ml tube and excess water was aspirated off. 600 μl of 60% isopropanol was added to fix animals and centrifugated at 1,200 relative centrifugal force (rcf) to pellet worms. The supernatant was removed and 600 μl of Oil Red O solution was added to each tube with pelleted worms. The Oil Red O solution was made using 0.5 g Oil Red O (Sigma, Cat # O0625) in 100 ml of 100% isopropanol, filtered through a 0.20 μm PVDF filter and allowed to equilibrate overnight with agitation at room temperature. Tubes were placed in a wet chamber and worms were stained for six hours at 25°C. After staining, worms were centrifugated at 1,200 rcf, washed twice, and re-suspended in 0.01% Triton X-100 in S-buffer. Worms were imaged on a 2% agar pad using a Leica DMI 3000-B inverted microscope coupled to a Leica DFC295 color camera. Oil-Red intensity was quantified using the Image J software (NIH). Pixel intensity was measured in the green color channel of the images. The region of the intestine measured on each animal was from the anterior part of the intestine (first cell) to region of the intestine in the mid-body at the same AP location as the vulva. Each worm was analyzed using an equivalently sized window.

Fixative Nile Red staining was performed on transgenic and non-transgenic worms as described [108]. Briefly, well-fed worms were age-synchronized by the bleaching method, and 500-1,000 L4-staged worms were washed from NGM plates seeded with *E. coli* OP50 using PBS containing 0.01% Triton X-100. Worms were allowed to settle by gravity and washed once with PBS. Excess PBS was removed and 200 μl of 40% isopropanol was added to fix animals for 3 minutes. Next, the supernatant was removed, and 150 μl of a Nile Red (Sigma, Cat # 19123) solution in isopropanol was added to the fixed animals and allowed to stain for 30 min in the dark with agitation. After staining, worms were allowed to settle and washed once with 1x M9 buffer and kept in the dark at 4°C before visualization. Stained worms were mounted on 2% agar pads and imaged on a Leica DM5500 Nomarski microscope equipped with a Hamamatsu Orca II camera.

Triglyceride (TAG) levels were determined with a Triglyceride assay kit (Biovision, Cat # K622). Worms were age-synchronized by the bleaching method and grown at 20°C until the L4 larval stage on NGM plates seeded with *E. coli* OP50. Worms were collected and washed with S-basal solution. A 5% Triton X-100 solution with 1x protease inhibitors (Roche Complete Mini, EDTA free) was added 1:1 to a 50 μl worm pellet, and worms were sonicated with a water bath sonicator (Branson). Lipids were dissolved twice by heating the lysate to 90°C for 5 min followed by vortexing. Following centrifugation, the supernatant was used to determine the total TAG levels according to the manufacturers protocol. TAG concentrations were normalized to the total protein content as determined by a Micro BCA protein assay kit (ThermoFisher, Cat # 23235). Each assay was done in triplicate and the average TAG level (μg TAG/ μg protein) was calculated.

### Assessment of lipid droplet morphology

The number and size of lipid droplets was measured as reported [109] in wild-type and *kin-29* null mutants expressing the transgenic DHS-3::GFP marker. Briefly, worms were collected at the L4 larval stage and imaged using a Leica SP8 confocal microscope with LAS software. Images were taken with a 63x objective. The anterior four intestinal cells were imaged, and the diameter and number of all visible droplets in a 50 μm^2^ area were measured using Image J version 1.51h (NIH) software [110].

### Assessment of food-leaving behavior

Food-leaving behavior was monitored by video recording 5 to 7 freely-moving young adult worms at 18°C over a 12 hour period. The area explored was quantified on a 5.5 cm diameter NGM agar plate freshly seeded with 5 μl *E. coli* OP50 that was grown to stationary phase on the night before seeding the plate. The bacteria formed a circle of 0.6 cm diameter in the middle of the plate. Movies were taken on a custom-built imaging system and worm tracking software (Volumetry, version 8.a) [111] using a USB 2.0 monochrome machine vision cameras (Point Grey Research, CMLN-13SM-CS) equipped with a 12.5 mm focal length C-mount lenses (Fujinon, HF12.5HA-1B). All imaging was performed under dark field illumination using low angle red-light LED rings as a light source, such that worms appeared as white objects against a dark background. All cameras were kept at the same height above the plates and bacterial lawns of the same diameter were used in all experiments. Videos were recorded in uncompressed QuickTime format using StreamPix software (Norpix, Montreal Canada) by capturing images at a rate of 0.5 frames/s. Video files were then imported into Volumetry, and each frame was converted into an 8-bit grayscale image for subsequent analysis. To quantify food-leaving behavior, we generated a binary image containing only white pixels when the grayscale value was above a user defined threshold that approached the maximum (intensity) grayscale value (255). This binary image identified the worms in each frame. We then collapsed the resulting frames into a single image to visualize worm tracks outside of the bacterial lawn for each 12-hour video. Worm track images were imported into the ImageJ software (NIH) and the average number of pixels representing worm tracks outside of the bacterial lawn was quantified

### Measurements of ATP levels

ATP levels in whole worms were determined as described [49]. For DTS experiments, ∼6,000-7000 worms were age-synchronized using the double-bleaching method [112], transferred to NGM agar surface (10 cm diameter) that was fully covered with a lawn *E. coli* OP50 and grown at 20°C. L1 animals were washed off the agar surface using a pipette filled with 5 ml of M9 buffer. The worm and bacterial suspension was allowed to settle through a 5 μm nylon mesh filter, which passes bacteria but traps the worms. The worms trapped by the filter were then flash frozen in liquid N_2_ and stored at −80°C until analysis. Worms were collected every hour on the hour between 12 hours and 21 hours after feeding developmentally-arrested L1 animals. Because at 20°C, lethargus occurs between 16.5-18.5 hours, these sampling times including animals before, during, and after L1 lethargus. For SIS experiments, one-day old adult worms were exposed while on an agar surface without peptone in the presence of a thin layer of bacteria to 1500 J/m^2^ UVC irradiation, 254 nm, ultraviolet irradiation. Following irradiation, worm samples were collected hourly on the hour between time 0 and 5 hours after irradiation. 30-40 adults were collected in a 1.5 mL microfuge tube under stereomicroscopal observation using a platinum wire. The samples were flash frozen in liquid N_2_ and stored at −80°C until analysis. For time-course experiments, sleep was identified by measuring the fraction of non-pumping L1 worms for DTS, and the minutes per hour of body movement quiescence for SIS. For measuring ATP levels in L4 animals, worms were grown until the mid L4 larval stage, and 50 animals per sample were collected in a 1.5 mL microfuge tube, flash frozen in liquid N_2_ and stored at −80°C until analysis.

All samples for ATP determination were treated identically. Following collection off the agar surface using water and a nylon mesh into 1.5 mL microfuge tubes, the worms were flash frozen in liquid nitrogen within 8-12 min of preparation time. In preliminary experiments we found that, while there was some time-dependent degradation of ATP in the first five minutes, the levels change minimally within the time window (8-12 minutes) of collection (**Figure S1A**). Samples of frozen worms were immersed in boiling water for 15 min and then placed on ice for 5 min. ATP was quantified in supernatants of worm solutions using an ATP Determination Kit (Molecular Probes, Cat # A22066) and a microplate reader (Synergy HT, Biotek) capable of luminescence measurements according to the manufacturers protocols. ATP concentrations were normalized to total protein content as determined by a Micro BCA protein assay kit (ThermoFisher, Cat # 23235). ATP was measured in technical triplicates, and the average ATP concentration per μg protein was calculated per biological sample with at least 3 biological experiments for each time point unless indicated otherwise.

### Measurement of luminescence in live animals

Luminescence was measured as previously described [53]. We used a Synergy HT microplate reader (Biotek) using a 590/35 nm emission filter. Black with clear flat bottoms microplates (Corning) were used by placing ∼20 worms of the strain PE254, which carry the *feIs4[Psur-5::luciferase::GFP*] transgene (PE254) in a well in 100 μl of M9 buffer. 50 μl of luminescence buffer (phosphate buffer pH 6.5, 0.1 mM D-luciferin (ThermoFisher, Cat # L2916), 1% DMSO and 0.05% Triton as final concentrations) was added to each well for a total volume of 150 μl. Luminescence of each well was read 3 min after adding luciferin. During incubation, with luciferin the microplates were shaken at a medium setting. Background measurements of luminescence were subtracted from readings. Luminescence readings were normalized to green fluorescent protein fluorescence, which was measured using a 528/20 nm emission filter.

### Measurement of p-AMPK levels

Activated phosphorylated AMPK levels (p-AMPK) were measured as previously described [113]. Worm samples for each genotype, condition, and time-point were prepared by removing the supernatant. Pelleted worms were mixed with 1 volumes of 2× sample loading buffer (200 mm Tris-Cl, pH 8.0, 500 mm NaCl, 0.1 mm EDTA, 0.1% Triton X-100, and 0.4 mm phenylmethylsulfonyl fluoride) and boiled for 10 min by immersion in a water bath. Worm lysates were electrophoresed on a 4-20% pre-cast SDS-polyacrylamide gel (Mini-Protean TGX Gels, Biorad) and electroblotted onto a nitrocellulose membrane (Trans-Blot Turbo Transfer Pack, Biorad) using a Trans-Turbo Blot transfer system (Biorad). The membrane was incubated in a blocking solution containing phospho-AMPKα Thr 172 (Cell Signaling Technologies, Cat # 2535S, 1:1000 dilution) or β-actin (Millipore, Cat # MAB1501R, 1:3000 dilution) and rocked at 4°C overnight, followed by incubation with anti-mouse (Invitrogen, Cat # 7076S, 1:5000 dilution) or anti-rabbit horseradish peroxidase antibody (Jackson ImmunoResearch, Cat # 7074S, 1:5000 dilution) for 1 hour at room temperature. The ECL Western blotting system (Clarity™ Western ECL Substrate) was used to detect the secondary antibodies on the membrane. Luminescence of the blot was visualized and captured using the Chemidoc V3 Touch Western Imager for mini-gels (Biorad). ImageJ version 1.51h (NIH) [110] was used to quantify the intensity of p-AMPK and actin bands.

### Measurement of body size

Body-length measurements were carried out by acquiring digital images of adult worms 24 hours after the L4 larval molt at 100x magnification. The length of the worm was traced with short line segments using Leica LAS software (NIH), and the sum of the line lengths was calculated. The tail was not included in the measurements.

### Optogenetics

Optogenetic activation of RIS was conducted by exposing animals carrying the transgene *Paptf-1::ChR2::mkate2* to blue light using the GFP filter of a Leica stereomicroscope equipped with a Leica EL6000 light source. L4 animals were transferred to plates seeded with DA837 *E. coli* bacteria supplemented with either 100 mM *all-trans* retinal dissolved in EtOH or EtOH vehicle alone and incubated overnight in the dark at 20°C. While monitored at 5-12× objective lens, the pumping rate of young adult worms was counted for 10 seconds prior to exposure to blue light, 10 seconds while exposed to blue light, and 10 seconds after blue light exposure. ATR plates were utilized within a week of seeding with *E. coli* and were stored in the dark until use.

### Histamine mediated pharmacogenetics inhibition

To generate transgenic worms expressing histamine-gated chloride channels (HisCl) in the sleep-promoting RIS neuron (*Pflp-11::HisCl*), the coding region of HisCl [114] as well as sequences 3’ to the gene including an splice acceptor SL2 sequence, the coding region for mCherry, and the *unc-54* 3’UTR) was amplified from the pNP471 (*Prig-3::HisCl::SL2::mCherry*) plasmid [114] while the *flp-11* promoter (1.0 kb) [76] was amplified from genomic DNA using the polymerase chain reaction (PCR). These fragments were combined using overlap extension PCR [115] and the final PCR product was injected into N2 worms at a concentration of 50 ng/μL along with pCFJ90 (*Pmyo-2::mCherry*) (AddGene) at a concentration of 2 ng/μL as a transgenesis marker, and 1 kb DNA ladder (NEB) to bring the final concentration up to 150 ng/μL. Two transgenic lines were generated, NQ1208 and NQ1209 (**S1 Table**).

For sleep deprivation experiments, worms expressing *Pflp-11::HisCl* were exposed to plates containing 10 mM histamine hydrochloride (Sigma, Cat # H2750) immediately prior to experiments. For SIS experiments, worms were age-synchronized using the bleaching method [116] and grown on NGM agar plates seeded with either *E. coli* DA837 or OP50 until worms reached adulthood. One-day old adults were transferred individually onto the agar surface of individual wells of a WorMotel PDMS chip filled with either NGM agar supplemented with histamine dissolved in water (10 mM final concentration of histamine) or water vehicle. Within 15 min, worms were UV irradiated (1500 J/m^2^ UVC irradiation, 254 nm) while on the WorMotel chip, and movement quiescence (min) was recorded for ∼8-10 hours as described above. For DTS experiments, mid-L4 animals were transferred to the WorMotel chip with NGM agar supplemented with either histamine dissolved in water (10 mM) or water vehicle, and body movement quiescence (min) was recorded for ∼15-20 hours during L4 lethargus as described above.

### Perhexiline treatment

100 mM Perhexiline (PHX) (Sigma, Cat# SML0120) in DMSO solution was diluted to 1 mM using dimethyl sulfoxide (final DMSO concentration of 1%) and 100 μl was spotted on the agar surface of *E. coli* OP50 seeded plates. These plates were allowed to dry before worms were placed on them. For SIS experiments, L4 larvae were exposed for ∼12 hours to PHX. One-day old adults were transferred individually to WorMotel wells filled with NGM agar supplemented with either 1 mM PHX in 1% DMSO or 1% DMSO vehicle. Worms were UV irradiated (1500 J/m^2^ UVC irradiation, 254 nm) within 15 min of transfer, and movement quiescence was recorded for ∼8-10 hours as described above. For DTS experiments, late L2-staged larvae were exposed to plates containing PHX ∼12 hours prior to experiments. Early L4-staged worms were transferred individually to the WorMotel wells containing NGM agar supplemented with either 1 mM PHX in 1% DMSO or DMSO vehicle. Animals were then videorecorded for 15-20 hours, which were later analyzed to obtain body movements quiescence measurements as described above.

### Subcellular localization of KIN-29(WT)::GFP and KIN-29(S517A)::GFP

An asynchronous population of gravid adult worms were treated with alkaline bleach and the progeny was allowed to enter the L1 diapause stage for 12 hours before resuming development by feeding them. For each transgenic line (PY5790 and NQ1241) and time point, between 1,500-2,500 worms were plated on 60 mm NGM agar plates seeded with bacteria. Worms were collected in 40% isopropanol and fixed for 1 min at room temperature on a nutator. Worms were sedimented by centrifugation at 800 rfc for 1 min. The supernatant was removed and 200 μl of 4,6-diamidino-2-phenylindole (DAPI) staining solution (2 ng/μl in PBST) was added to the worm pellet. Worms were allowed to stain for 2 min in the dark with nutation, and pelleted by centrifugation at 1200 rfc for 1 min. The supernatant was removed and worms were washed 1× with PBST to remove excess DAPI. Worm were transferred to an agar pad and image z-stacks (between 10 images with a step size of 0.7 μm at 60x magnification) are captured with a Leica SP8 confocal microscope in a sequential fashion (altered between GFP to DAPI). To quantify subcellular localization of KIN-29::GFP, an experimentor was blinded to the genotype/condition of the worm, and each worm was scored on a scale of 0-3, where 0 denotes fully cytoplasmic and 3 denotes fully nuclear location of the GFP.

### Statistical analysis

Graphpad Prism 7 software was used for data analysis. Statistical comparisons were made using the two-tailed Mann-Whitney *t*-test (unpaired, non-parametric) or two-way ANOVA with corrections for multiple hypothesis testing unless indicated otherwise. Specific statistical tests are described in the figure legends.

## ACKNOWLEDGEMENTS

We thank Victoria Chen for performing fat staining during lethargus, Michael Iannacone for constructing the *ceh-17; aptf-1* double mutant strain, Mark Nessel for constructing the *kin-29;hs:EGF* and *kin-29;hs:FLP-13* strains, members of the Raizen and van der Linden labs, Amita Sehgal, and Matthew Kayser for discussions and comments on this manuscript; the *Caenorhabditis* Genetics Center (CGC) and Piali Sengupta for strains; Matt Churgin and the lab of Christopher Fang-Yen for assistance with the WorMotel device; the Cellular and Molecular Imaging Core Facility of the COBRE Integrative Neuroscience Center for providing equipment necessary for Western blotting.

## SUPPLEMENTARY FIGURE LEGENDS

**S1 Fig. Total ATP and p-AMPK levels in *aak-2* and *pink-1* mutants, as well as movement quiescence and ATP levels during sleep deprivation.**

**(A)** ATP extinction curve of wild-type L4 animals (see **Material and Methods**) as a function of time after sample extraction. **(B)** Quantification of phosphorylated AMPK (p-AMPK) levels in L4 animals of *aak-2* null mutants with representative Western blots where the intensity of the bands represents p-AMPK (top panel) and β-actin (lower panel) as a loading control. Data are normalized to wild-type and represented as the mean ± SEM of 2 experiments. *** p<0.001. A sequence alignment of AMPK proteins:phosphorylation site. AAK-2, the worm homolog of the AMPKα subunit is regulated by phosphorylation of Threonine-243 (purple), which corresponds to the Threonine-172 of human AMPKα. **(C and D)** Levels of total body ATP (nM ATP/μg protein) (C) and phosphorylated AMPK (p-AMPK relative to actin loading control) (D) measured in L4 animals of wild-type controls and *pink-1* mutants. Data are normalized to wild-type controls and represents as the mean ± SEM of 6 experiments for ATP, and 2-3 experiments for p-AMPK. * p<0.05 by a two-tailed unpaired *t*-test, ** p<0.01. Representative Western Blots are shown of wild-type and *pink-1* mutants where the intensity of the bands represents p-AMPK (top panel) and β-actin (lower panel) as a loading control. **(E and F)** Pharmacogenetic silencing of RIS neurons results in a reduction in body movement quiescence during L4 lethargus/DTS and after UVC exposure/SIS in wild-type animals expressing the *Pflp-11::HisCl* transgene (two independent transgenic lines, NQ1208 and NQ1209) in the presence of 10 mM Histamine (+His), and in the absence of histamine (-His). The *flp-11* promoter is expressed in RIS. Left graphs: Time-course with minutes of quiescence in 10-min bins during L4 lethargus/DTS and minutes of quiescence in 1-hr bins. Statistical comparisons were performed with a 2-way ANOVA using time and genotype as factors, followed by post-hoc pairwise comparisons at each time point to obtain nominal p values, which were subjected to a Bonferroni correction for multiple comparisons. ***, ** and * indicates corrected p values that are different from transgenic animals (-His) at p<0.001, p<0.01 and p<0.05, respectively. Right graphs: Total minutes of quiescence during L4 lethargus/DTS and quiescence during the first 4 hours after UVC irradiation (1,500 J/m^2^) determined from the time-course data. Data are represented as mean ± SEM. * p<0.05, ** p<0.01, *** p<0.001.

**S2 Fig. Quiescence, total ATP and p-AMPK levels in *kin-29* mutants.**

**(A)** SIK phylogeny tree. The bootstrap values of each branch are denoted in red. All alignments were performed using the MUSCLE alignment algorithm and the tree was generated with the Phylogeny.fr program (http://phylogeny.lirmm.fr/phylo_cgi/index.cgi) [117] Sequences (UniProtKB) used were as follows: SIK3 *Drosophila* (Q4QQA7), SIK2 *Drosophila* (O77268), SIK1 mouse (Q60670), SIK2 mouse (Q8CFH6), SIK3 mouse (Q6P4S6), SIK1 human (P57059), SIK2 human (Q9H0K1), SIK3 human (Q9Y2K2) and KIN-29 *C. elegans* (Q21017). **(B)** Heatmap of the fraction of movement quiescence of 7 wild-type and 7 *kin-29* null mutants during L4 lethargus/DTS. Each row in the heatmap represents an individual animal recorded by video over a ∼20-hour period. The time of each worm’s record was adjusted to align to the start of L4 lethargus quiescence. See **Materials and Methods**. **(C)** Fraction of quiescence of a wild-type and *kin-29* mutant individual animal during L4 lethargus/DTS. The time of the two worms’ records were adjusted to align to the start of the L4 lethargus quiescence. **(D)** Feeding rate of wild-type and *kin-29* mutant animals after UVC irradiation (1,500 J/m^2^). Horizontal line denotes the mean ± SEM (n>10 animals). *** p<0.01, Student’s two tailed *t* test. **(E and F)** Body movement quiescence (E) is reduced and feeding rate (F) is increased after heat shock/SIS is reduced in *kin-29* mutant animals in comparison to wild-type. Adult animals were heat-shocked at 35°C for 30 minutes (see **Material and Methods**). Graphs show the mean ± SEM of n=19-23 animals for movement quiescence and n=9-15 animals for feeding rate. * p<0.05, ** p<0.01, *** p<0.001, Student’s two tailed *t* test. **(G)** Bioluminescence in wild-type and *kin-29* mutant animals carrying the *sur-5::luciferase::gfp* reporter. Luminescence was normalized to *gfp* expression emanating from the same transgene. Data are represented as the mean ± SD of 6-7 replicates. ** p<0.01, Student’s two tailed *t* test. **(H)** Quantification and representative Western Blots of p-AMPK in L4 larvae of wild-type and *kin-29* mutants. β-actin (lower panel) is a loading control. Data are normalized to wild-type and represented as the mean ± SEM of 3 experiments. ** p<0.01, Student’s two tailed *t* test. **(I)** Levels of total body ATP in wild-type and *kin-29* mutant animals at 0 and 2 hours after induction of SIS by UVC exposure. Data were normalized to wild-type controls immediately before UVC exposure (0 hours) and are represented as the mean ± SEM of 4-6 experiments, ** p<0.01, Student’s two tailed *t* test.

**S3 Fig. Fat content, food-uptake and food-leaving behavior in *kin-29* mutants.**

**(A)** Food-leaving behavior measured in wild-type controls, *kin-29*, and *eat-2* mutant animals. *kin-29* mutants have increased food leaving behavior similar to *eat-2* feeding defective mutants Uptake was measured 15-min after exposure to fluorescent microsphere beads of 1.0 μm diameter. Data is represented as the mean ± SEM (n=10-13 animals). *** p<0.001, Student’s two tailed *t* test. **(B)** Fat content measured with fixative Oil-Red O staining for wild-type and *kin-29* mutants. Data are represented as the average percentage of total body fat in wild-type controls ± SEM (n=12 animals for each genotype and developmental stage). *** and * indicate values that are different from wild-type at p<0.001 and p<0.05, respectively, Student’s two tailed *t* test. Representative images are shown of animals fixed and stained with Oil Red O at different developmental stages from L1 larvae to young adults of wild-type and *kin-29* mutants. **(C)** Lipid droplet morphology of *kin-29* mutants. Animals mutant for *kin-29* result in an increased lipid droplet number and size. Expression of *kin-29* in *odr-4*-expressed neurons restores the increased lipid droplet number of *kin-29* mutants. Lipid droplet number and size is quantified within a 50 or 18 μm^2^ area, respectively, in the anterior intestine. ** p<0.01, *** p<0.001, Student’s two tailed *t* test. Middle panel: The distribution and average size of lipid droplets in wild-type and *kin-29* mutants. Lower panel: Representative images are shown of animals expressing DHS-3::GFP in wild-type and *kin-29* mutants. DHS-3::GFP stains the membrane of lipid droplets in gut cells. **(D)** Microsphere uptake measured in wild-type controls, *kin-29*, and *eat-2* mutant animals. Animals mutant for *kin-29* have normal food uptake unlike *eat-2* feeding defective mutants Representative images are shown of single adult wild-type, *kin-29*, and *eat-2* mutant animals that accumulated 1.0 μm fluorescent microsphere beads for 15 min at 20°C. The arrows indicate the terminal bulbs of the pharynxes.

**S4 Fig. Rescue of the reduced DTS and SIS of *kin-29* mutants by ATGL-1 overexpression.**

**(A)** ATGL-1 overexpression (OE) restores in part the defect in DTS body movement quiescence of *kin-29* L4 null mutants. Data are represented as a moving window of the fraction of a 10-min time interval spent quiescent of n=6 animals for each trace. The x-axis represents hours from the start of recording in the late fourth (L4) larval stage. The data from individual worms was aligned such that the start of lethargus quiescence occurred simultaneously. **(B)** Time-course of minutes of quiescence in 1-hr bins after UVC irradiation (1,500 J/m^2^). Data are represented as the mean ± SEM (n=8-10 animals for each genotype). Statistical comparisons were performed with a 2-way ANOVA using time and genotype as factors, followed by post-hoc pairwise comparisons at each time point to obtain nominal p values, which were subjected to a Bonferroni correction for multiple comparisons. ** indicates corrected p values that are different from wild-type at p<0.01.

**S5 Fig. Perhexiline treatment reduces DTS and SIS in wild-type animals**

**(A)** Schematic showing that the carnitine palmitoyltransferase (CPT) inhibitor perhexiline (PHX) blocks beta oxidation of fatty acids in mitochondria to promote the accumulation of lipids. **(B)** Perhexiline (1 mM) reduces body movement quiescence during L4 lethargus/DTS in wild-type animals in comparison to vehicle controls (-PHX). Data are represented as a moving window of the fraction of a 10-min time interval spent quiescent of n=9 animals for the PHX(-) trace and n=11 animals for the PHX(+) trace. The x-axis represents hours from the start of recording in the late fourth (L4) larval stage. The data from individual worms was aligned such that the start of lethargus quiescence occurred simultaneously. **(C-D)** Time-course of minutes of quiescence in 1-hr bins (C) and feeding rate (in pumps per 10 sec) (D) after UVC irradiation (1,500 J/m^2^) in the absence (-) and presence (+) of perhexiline (PHX). Data are represented as the mean ± SEM with n=19 animals for the PHX(-) condition and n=28 animals for the PHX(+) condition. Statistical comparisons were performed with a 2-way ANOVA using time and PHX conditions as factors, followed by post-hoc pairwise comparisons at each time point to obtain nominal p values, which were subjected to a Bonferroni correction for multiple comparisons. ** and *** indicates corrected p values that are different from wild-type at p<0.01, and p<0.001, respectively.

**S6 Fig. Rescue of *kin-29* mutant phenotypes by expressing *kin-29* in a subpopulation of sensory neuron types.**

**(A)** Representative images of animals fixed and stained with Nile Red (left images) food-leaving behavior (black/white image) of adult animals expressing the *Podr-4::kin-29* or *Pges-1::kin-29* transgene. *odr-4*, chemosensory promoter. *ges-1*, intestinal promoter. Frames from a 12-hour video were collapsed in a single image for food-leaving behavior. **(B)** Total ATP levels (nM ATP/μg protein) in wild-type and *kin-29* null mutant animals expressing the *Podr-4::kin-29* transgene measured before, during and after L1 lethargus/DTS. Data are normalized to the average value of the wild-type time-course. The graphs show the mean ± SEM of 2 experiments for wild-type, and 3 experiments for *kin-29* animals that carry the extra-chromosomal *Podr-4::kin-29* transgene (PY5791). Of note, PY5791 includes about 20% non-transgenic *kin-29* animals. **(C)** Time-course of minutes of quiescence in 1-hr bins after UVC irradiation (1,500 J/m^2^). Data are represented as mean ± SEM (n=8-10 animals). Statistical comparisons were performed with a 2-way ANOVA using time and genotype as factors, followed by post-hoc pairwise comparisons at each time point to obtain nominal p values, which were subjected to a Bonferroni correction for multiple comparisons. *** and * indicates corrected p values that are different from *kin-29* mutants at p<0.001 and p<0.05, respectively.

**S7 Fig. Rescue of *kin-29* mutant phenotypes by expressing *kin-29* in individual or a subpopulation of sensory neuron types.**

**(A)** Schematic showing the transgenic rescue strategy used to restore *kin-29* expression in a subset of *odr-4*-expressing sensory neurons (left) and in individual neurons (right). **(B)** Expression and localization of a *srh-56* promoter (∼1.5 kb) fusion with GFP. A black and white image shows *Psrh-56::gfp* fluorescence in ASK, ASH and ASJ neurons plus other non-neuronal cells. **(C)** Representative images of a single L4 larvae fixed and stained with Oil Red O of wild-type and *kin-29* null mutant animals with or without the *Psrh-56::kin-29* transgene. **(D)** Time-course of minutes of quiescence in 1-hr bins after UVC irradiation (1,500 J/m^2^) of wild-type and *kin-29* null mutants with or without the *Podr-3::kin-29* transgene. Data are represented as the mean ± SEM (n=14-15 animals). Statistical comparisons were performed by an unpaired multiple comparison *t*-test with Holm-Sidak correction. *** and * indicate corrected p values that are different from *kin-29* mutants at p<0.001 and p<0.05, respectively. **(E)** Fraction of quiescence of *kin-29* null mutants expressing *kin-29* under control of the *odr-3* (AWA, AWB, AWC, ADF, ASH), *gpa-4* (ASI), *srh-142* (ADF) and *sre-1* (ADL) promoter. Data are represented in a 10-min time interval of at least n=4 animals for each sleep trace. The x-axis represents hours from the start of recording in the late fourth (L4) larval stage.

**S8 Fig. RIS is required for SIS, and *kin-29* mutants do not attenuate the quiescence of FLP-13 overexpression.**

**(A and B)** Body movement quiescence (A) and feeding rate (B) of *kin-29* null mutants after induction of FLP-13 overexpression (OE) (n>14 animals for each genotype). To induce FLP-13 overexpression, adult animals expressing a *Phsp-16.2::FLP-13* transgene were heat shocked for 30 min (see **Material and Methods**). Data are represented as the mean ± SEM. *** p<0.001, Student’s two tailed *t* test. **(C to F)** Body movement quiescence and feeding rate following either UVC irradiation 1,500 J/m^2^ (C and D) or heat shock 35°C for 30 minutes (E and F). Data are represented as mean ± SEM (n>10 animals for each genotype). For the time-course experiments, statistical comparisons were performed with a 2-way ANOVA using time and genotype as factors, followed by post-hoc pairwise comparisons at each time point to obtain nominal p values, which were subjected to a Bonferroni correction for multiple comparisons. *** p<0.001, ** p<0.01, and * p<0.05.

**S9 Fig. KIN-29(S517A) mutant rescues the small body size phenotype of *kin-29*.**

**(A)** Sequence alignment of SIK3 proteins (human Q9Y2K2, mouse Q6P4S6, *Drosophila* Q4QQA7, *C. elegans* Q21017). The conserved PKA phosphorylation site Serine is shown in blue. **(B)** KIN-29(S517A) remains cytosolic after heat shock. Shown is KIN-29(WT)::GFP in the nucleus (arrow) of an *odr-4*-expressed neuron after heat shock (left image). KIN-29(S517A)::GFP remains in cytoplasmic after heat shock (right image). **(C)** KIN-29(S517A) rescues the small body size of *kin-29* null mutants. Shown is the relative body length of adult animals (n=8-13) of the indicated genotypes. Data are represented as the mean ± SEM. *** p<0.001, Student’s two tailed *t* test.

**S10 Fig. Mutations in *hda-4* restore *kin-29* mutant phenotypes with the exception of fat stores.**

**(A)** Time-course of minutes of quiescence in 1-hr bins after UVC irradiation (1,500 J/m^2^) of single and double mutants between *kin-29* and *hda-4* compared to wild-type. Data are represented as the mean ± SEM (n>10 animals). Statistical comparisons were performed with a 2-way ANOVA using time and genotype as factors, followed by post-hoc pairwise comparisons at each time point to obtain nominal p values, which were subjected to a Bonferroni correction for multiple comparisons. ***, ** and * indicates corrected p values that are different from wild-type at p<0.001, p<0.01 and p<0.05, respectively. **(B)** Time-course of minutes of quiescence in 1-hr bins after UVC irradiation (1,500 J/m^2^) of animals expressing *hda-4* under the control of its own promoter and under the control of the *odr-4* chemosensory neuron specific promoter. Data are represented as the mean ± SEM (n>10 animals). Statistical comparisons were performed with a 2-way ANOVA using time and genotype as factors, followed by post-hoc pairwise comparisons at each time point to obtain nominal p values, which were subjected to a Bonferroni correction for multiple comparisons *** and ** indicates corrected p values that are different from wild-type at p<0.001, and p<0.01, respectively. **(C)** Representative images of animals fixed and stained with Oil Red O (left images) and food-leaving behavior (black/white image) of wild-type, single and double mutant animals of *kin-29* and *hda-4*. Frames from a 12-hour video were collapsed in a single image for food-leaving behavior. **(D)** Total levels of ATP (nM ATP/μg protein) and phosphorylated AMPK (p-AMPK relative to actin loading control) in wild-type and *kin-29 hda-4* double mutants measured before, during and after L1 lethargus/DTS. ATP levels in wild-type worms are not significantly different from *kin-29hda-4* double mutants (2-way ANOVA multiple comparison with Bonferroni correction). Shown is a representative time-course of ATP and p-AMPK measured from the same samples. Data are normalized to the average value of the wild-type time-course. Graphs show the mean ± SD for wild-type and *kin-29hda-4* doubles. A Western Blot is shown below the graphs for *kin-29hda-4* doubles with p-AMPK (top panel) and β-actin (lower panel) as a loading control. **(E)** Quantification and representative Western Blots of p-AMPK in L4 larvae of wild-type and single and double mutants of *kin-29* and *hda-4*. β-actin (lower panel of the blot) is used as a loading control. Data are normalized to wild-type. ** and * indicate values that are different from wild-type at p<0.01 and p<0.05, respectively, Student’s two tailed *t* test.

